# Network- and Measure-Specific Mid-Term Reliability of Multi-Echo Resting-State Functional Magnetic Resonance Imaging on a Compact 3 Tesla Scanner

**DOI:** 10.64898/2026.08.07.743542

**Authors:** Daehun Kang, Kirk M Welker, Dora Hermes, Matt A Bernstein, John Huston, Yunhong Shu

## Abstract

1.

**Introduction:** Understanding mid-term test-retest reliability and within-subject variability is important for interpreting changes observed in longitudinal and intervention studies. The reliability of resting-state functional magnetic resonance imaging (rs-fMRI) is known to vary across measures and brain regions. However, how reliability differs across functional networks and connectivity-and amplitude-based measures, and whether multi-echo acquisition and processing modify these patterns, remain incompletely characterized.

**Methods:** Twenty-two healthy volunteers underwent two rs-fMRI sessions 15.7 ± 4.0 days apart on a Compact 3T scanner. Multi-echo, middle-echo, and independently acquired single-echo datasets were compared, with multi-echo independent component analysis additionally evaluated as a denoising approach. Functional connectivity (FC) and three amplitude-based measures were evaluated using the Schaefer 400 parcellation. Reliability was systematically assessed using intraclass correlation coefficient (ICC), within-subject standard deviation (wSD), and systematic bias at edge or regional, and network levels.

**Results:** Acquisition-dependent differences in reliability were generally modest. Multi-echo acquisition and processing increased functional connectivity strength and the magnitude of amplitude-based measures and improved inferior cortical coverage, but these enhancements did not consistently translate into substantially higher ICC or lower wSD. In contrast, reliability showed clear network-dependent differences. FC reliability varied markedly across network pairs and was not explained by connectivity strength alone; pairs involving the default mode and control networks generally showed more favorable profiles than several somatomotor and visual network pairs. Fractional amplitude of low-frequency fluctuations (fALFF) also showed network-dependent reliability, with the most favorable regional reproducibility observed in the default mode and control networks and lower reproducibility in the somatomotor and visual networks.

**Conclusion:** These findings provide practical mid-term reliability benchmarks for rs-fMRI on a Compact 3T scanner and show that measurement stability varies more clearly across measures and functional networks than across acquisition approaches.

**Key points:**

- Mid-term test-retest reliability varied more clearly across resting-state measures and functional networks than across acquisition and processing approaches.
- Multi-echo acquisition and processing enhanced functional connectivity strength, amplitude-based signal magnitude, and inferior cortical coverage but did not consistently improve reliability.
- Functional connectivity strength and fractional amplitude of low-frequency fluctuations showed distinct network-specific reliability profiles, with more favorable reproducibility in default mode and control networks than in several somatomotor and visual networks.

## 2. Introduction

Understanding the reliability and variability of resting-state fMRI measures, including functional connectivity and amplitude-based measures, is essential for longitudinal and intervention studies, because observed changes may reflect both meaningful biological or intervention-related effects and ordinary scan-to-scan variability (Zuo, Di Martino et al. 2010, Blautzik, Keeser et al. 2013, O’Connor, Potler et al. 2017, Conwell, von Reutern et al. 2018, Ning, Makris et al. 2019, Noble, Scheinost et al. 2019). This detection is particularly important when changes in resting-state fMRI measures occur without corresponding behavioral or clinical changes (Noble, Spann et al. 2017, Kang, Uchida et al. 2026). Even when biologically meaningful changes are observed, it remains important to determine whether resting-state fMRI measures primarily reflect stable trait-like brain characteristics or state-dependent variability at the time of scanning (Finn, Shen et al. 2015, Geerligs, Rubinov et al. 2015). To address this question, the present study used a mid-term test–retest design to evaluate measurement stability across independent scan sessions. The approximately 2-to 3-week interval was chosen to capture scan-to-scan and state-dependent variability while minimizing the likelihood of major biological change in healthy individuals. These considerations are particularly relevant when evaluating emerging acquisition platforms and sequence designs.

Reliability may also vary according to the resting-state fMRI measures being examined (Somandepalli, Kelly et al. 2015, Cahart, O’Daly et al. 2023). Functional connectivity characterizes temporal synchronization between brain regions (Biswal, Yetkin et al. 1995), whereas amplitude-based measures, including amplitude of low-frequency fluctuations (ALFF), fractional ALFF (fALFF), and mean-normalized ALFF (mALFF) (Yang, Long et al. 2007, Zang, He et al. 2007, Zou, Zhu et al. 2008), characterize the magnitude of spontaneous low-frequency BOLD fluctuations. Because these measures differ in their calculation, signal scaling, and sensitivity to physiological and non-neural fluctuations, they may exhibit distinct test-retest reliability profiles (Zuo, Di Martino et al. 2010, Noble, Scheinost et al. 2019). Reliability may also vary across large-scale functional networks because of differences in network organization, connectivity strength, signal quality, anatomical location, and susceptibility to state-dependent modulation (Zuo, Kelly et al. 2010, Geerligs, Rubinov et al. 2015, Noble, Spann et al. 2017). Moreover, because edge-wise or ROI-wise ICC estimates can be noisy, network-level summaries may provide a more stable description of system-level reliability profiles (Mejia, Nebel et al. 2015, Pannunzi, Hindriks et al. 2017). Therefore, assessing reliability jointly across acquisition strategies, resting-state fMRI measures, and large-scale functional networks may provide a more informative characterization of reliability than relying on a single global reliability estimate.

The intraclass correlation coefficient (ICC) is widely used to assess test-retest reliability because it quantifies the extent to which inter-individual differences are preserved across sessions (McGraw and Wong 1996). However, ICC depends on both within-subject and between-subject variance and therefore does not fully characterize absolute scan-to-scan variability or systematic session effects (Chen, Xu et al. 2015, Noble, Scheinost et al. 2019). We therefore complemented ICC with the within-subject standard deviation (wSD), which quantifies absolute within-subject variability, and test-retest bias, which measures systematic shifts between sessions (Bland and Altman 1986). Together, these metrics provide complementary assessments of relative reliability, absolute variability, and systematic test–retest change.

Recent advances in high-performance head-only MRI systems have introduced hardware configurations that differ from those of conventional whole-body scanners, including Compact 3T, SIGNA MAGNUS system, Connectome 2.0, Compact 7T and NexGen 7T systems, (Foo, Laskaris et al. 2018, Foo, Tan et al. 2020, Feinberg, Beckett et al. 2023, Ramos-Llorden, Lee et al. 2026, Wu, Ricci et al. 2026), which could affect EPI distortion, temporal sampling, signal stability, and the sensitivity of resting-state fMRI measures (Kang, Jo et al. 2020). This study was performed on a compact 3T MRI system (C3T), a lightweight, low-cryogen, head-only scanner equipped with a high-performance gradient system with a peak slew rate of 700 T/m/s and a peak gradient amplitude of 80 mT/m on all gradient axes simultaneously (Foo, Laskaris et al. 2018). The gradient performance enables shorter echo train duration and allowed three-echo EPI acquisition to be implemented with ∼ 1 s TR and 2.4 mm isotropic resolution (Kang, In et al. 2023). Although multi-echo acquisition and echo combination improves BOLD sensitivity and mitigate susceptibility-related signal loss, it remains unclear whether the signal and sensitivity advantages translate into improved test–retest reliability of resting-state fMRI measures. The reliability of optimally combined multi-echo, middle-echo–derived, independently acquired single-echo, and multi-echo ICA-denoised data has not been systematically evaluated on this compact 3T platform.

In this study, we evaluated the mid-term test–retest reliability of resting-state fMRI metrics acquired on a compact 3T MRI scanner using multi-echo EPI, a middle-echo dataset derived from the multi-echo acquisition and independently acquired single-echo EPI acquisitions. We examined functional connectivity and amplitude-based measures, including ALFF, fALFF, and mALFF, at ROI, edge, and network levels. Specifically, we investigated whether test–retest reliability differed across acquisition strategies, whether commonly used resting-state metrics exhibited distinct stability profiles, and whether reliability varied systematically across large-scale functional networks. By jointly assessing relative reliability, absolute within-subject variability, and systematic test–retest shift using ICC, wSD, and bias, respectively, this study provides practical benchmarks for interpreting resting-state fMRI changes over clinically relevant mid-term intervals.

## 3. Materials and Methods

### 3.1. Data acquisition

Following a study protocol approved by the Institutional Review Board, 22 healthy volunteers (15 females / 7 males; 40.8 ± 13.7 years old) participated in this study after written informed consent was obtained. All subjects were scanned on two separate sessions with inter-session intervals of 15.7 ± 4.0 days.

All MRI scans were conducted with the Compact 3T MRI scanner as a technology demonstrator (Foo, Laskaris et al. 2018), using a 32-channel brain coil (Nova Medical, Inc., Wilmington, MA, USA). The fMRI experiments were performed using a multi-echo multi-band gradient-echo EPI sequence with blipped-controlled aliasing in parallel imaging technique (Setsompop, Gagoski et al. 2012, Cohen, Yang et al. 2021). Resting-state fMRI datasets were acquired using both conventional single-echo EPI and multi-echo EPI acquisitions. For the multi-echo acquisition, three echoes (TE = 11.7, 29.7, and 47.7 ms) were acquired. In addition to T2*-weighted multi-echo combination (MEPI), the middle echo (MMID) (TE = 29.7 ms) was extracted and analyzed separately to enable comparison between multi-echo, middle-echo– derived, and conventional single-echo acquisitions (SEPI). A whole-brain fMRI protocol, including the following imaging parameters were applied in Table 1. The longer TR for the multi-echo acquisition was required to accommodate acquisition of three echoes while maintaining comparable spatial coverage and scan duration.

**Table 1.** Details of imaging parameters for single-and multi-echo EPI acquisitions.

| Imaging Parameter | Single-Echo EPI | Multi-Echo EPI |
| --- | --- | --- |
| Field of view (FOV, mm) | 224 × 224 |  |
| Matrix size | 94 × 94 |  |
| In-plane resolution (mm) | 2.38 × 2.38 |  |
| k-space partial Fourier factor | No partial Fourier |  |
| Slice thickness (mm) | 2.4 |  |
| Number of slices | 52 |  |
| Inter-slice gap (mm) | 0.3 |  |
| Inferior-to-superior coverage (mm) | 140.4 |  |
| In-plane acceleration factor | 2 |  |
| Multi-band factor | 4 |  |
| Echo spacing (μs) | 352 |  |
| Receiver bandwidth (Hz) | ±250 k |  |
| Echo time (TE, ms) | 30 | 11.7, 29.7, 47.7 |
| Repetition time (TR, ms) | 700 | 940 |
| RF flip angle (FA, °) <sup>1</sup> | 52 | 59 |
| Number of volumes per scan | 886 | 660 |
| Scan duration (mm:ss) | 10:21 | 10:20 |
| Note | - | - |

<sup>1</sup> FAs were applied based on the calculated Ernst angles assuming T<sub>1</sub> of 1.4 s in gray matter at 3T.

During the resting-state fMRI scans, all subjects were instructed to remain still and fix their gaze on the center of a mirror mounted on the head coil. Independent measures of physiological variables (cardiac and respiration) were recorded. At each scan session, participants completed a brief behavioral and physiological questionnaire assessing sleep duration and quality, current restedness, fluid and caffeine intake, fed/fasted state, and physical discomfort (Supplementary Material 1), developed with reference to the hydration, caffeine, and sleep questionnaire described in previous work (Gorgolewski, Mendes et al. 2015). Scan acquisition time was extracted from the DICOM metadata.

### 3.2. Data Preprocessing

The preprocessing procedures for RS-fMRI datasets were all carried out using AFNI (Analysis of Functional NeuroImages, https://afni.nimh.nih.gov/)’s suite of programs (Cox 1996). For preprocessing and artifact reduction of the RS-fMRI data, the following steps were performed in sequence: truncation by initial ten volumes, de-spiking (‘3dDespike’ in AFNI), physiological noise elimination including cardiac and respiratory artifacts (‘RETROICOR’ and ‘RVT’) (Glover, Li et al. 2000, Birn, Smith et al. 2008), slice acquisition timing correction (‘3dTshift’ in AFNI), time-series alignment, Legendre polynomial detrending and motion-and hardware-related linear regressions (Jo, Saad et al. 2010), in order. For the ME dataset, a T_2_*-weighted echo combination was performed between the time-series alignment and the detrending and the regressions (Posse, Wiese et al. 1999, Poser, Versluis et al. 2006, Heunis, Breeuwer et al. 2021) by using AFNI (‘@compute_OC_weights’). All datasets were evaluated for artifacts due to abrupt head motion, passing the sudden motion detection of AFNI at the threshold level of 0.3 mm for the Euclidean L2 norm of motion displacement during each TR interval (Jo, Gotts et al. 2013). To maintain identical temporal sampling across acquisitions and sessions, volume censoring was not applied.

T1-weighted anatomical images were processed to segment brain structures using FreeSurfer (Reuter, Schmansky et al. 2012). The resulting anatomical segmentations were used for coregistration to individual EPI images and for projection onto standard cortical surfaces provided by AFNI and SUMA (Cox 1996). Regions of interest (ROIs) were defined using the 400-region cortical parcellation from the Schaefer atlas (Schaefer, Kong et al. 2018). The atlas was implemented on the AFNI standard surface model (https://afni.nimh.nih.gov/pub/dist/atlases/SchaeferYeo/) and subsequently transformed into individual subjects’ native volumetric space for analysis of EPI data. Each ROI was assigned to one of 17 intrinsic connectivity networks, enabling analyses at both the ROI and network levels for functional connectivity and spontaneous BOLD signal fluctuations. Within-network FC indicated the edges connecting ROIs within the same network, whereas between-network FC was the edges connecting ROIs to different networks. Network-pair FC strength and reliability metrics were summarized using a median value across constituent edges.

#### 3.2.1. Multi-echo ICA-based denoising

To benchmark the effects of multi-echo ICA-based denoising, the multi-echo EPI data were additionally processed using TE-Dependent ANAlysis (‘tedana’ v26.0.3) for multi-echo independent component analysis (ME-ICA) (Kundu, Inati et al. 2012, DuPre, Salo et al. 2021). For each run, the initial ten volumes were removed, followed by de-spiking (3dDespike in AFNI), slice acquisition timing correction (3dTshift in AFNI), and time-series alignment/motion correction within afni_proc.py using AFNI. The three preprocessed echo images were then used as inputs to the default tedana workflow. The denoised T2*-weighted combined output generated by tedana (desc-denoised_bold.nii.gz) was passed back to afni_proc.py for the remaining processing steps, including Legendre polynomial detrending and motion-related linear regression. The resulting residual time series was defined as the MICA dataset and underwent the same downstream processing and analysis steps as the other functional datasets, including atlas-based time-series extraction, FC and ALFF-variant estimation, and test–retest reliability analysis. MICA was included as an additional benchmarked processing strategy to characterize the effects of multi-echo ICA-based denoising on FC magnitude, ICC, within-subject standard deviation, and test–retest bias.

### 3.3. Resting-state fMRI measures

To estimate amplitude-based resting-state fMRI measures of ALFF, fractional ALFF (fALFF), and global mean ALFF (mALFF) (Yang, Long et al. 2007, Zang, He et al. 2007, Zou, Zhu et al. 2008), voxel-wise calculation was performed for with options of bandpass-filtering with a frequency range of 0.01 to 0.10 Hz and smoothed by a Gaussian kernel with a 4 mm full-width-at-half-maximum on volume dataset by ‘3dRSFC’ in AFNI, and ROI-based summary including an option of ‘non-zero mean’ was conducted with 400 ROIs by ‘3dROIstats’ in AFNI.

For functional connectivity (FC) estimation, the band-passed and spatial-smoothed preprocessed residual fMRI datasets were re-used to extract time-series for 400 ROIs. FC strength was estimated with Pearson correlation between the time-series by ‘@ROI_Corr_Mat’ in AFNI. The resulting correlation coefficients were Fisher z-transformed before subsequent FC strength and test–retest reliability analyses.

### 3.4. Test–retest reliability metrics and Statistical comparisons

Test–retest reliability was evaluated using both relative and absolute reliability metrics. Relative reliability was assessed using the intraclass correlation coefficient (ICC), calculated in MATLAB using the ICC.m function with the ‘A-1’ option (version 1.3.1.0) (McGraw and Wong 1996, Salarian 2016). This option estimates single-measure absolute agreement among repeated measurements, rather than consistency alone.

Absolute within-subject variability was quantified using the within-subject standard deviation (wSD). For each resting-state fMRI measure (x) such as FC, ALFF, fALFF, and mALFF, wSD was calculated as:

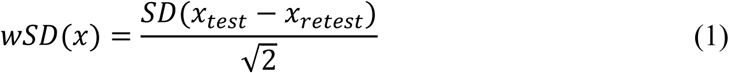

Systematic test–retest shift was quantified as the absolute bias:

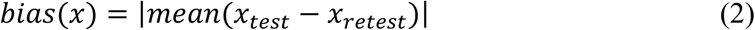

Because FC, ALFF, fALFF, and mALFF have different numerical scales, scale-normalized metrics (CVwSD and CVBias) were calculated to facilitate comparisons across metrics and acquisitions. CVwSD was defined as wSD divided by the absolute mean test–retest magnitude:

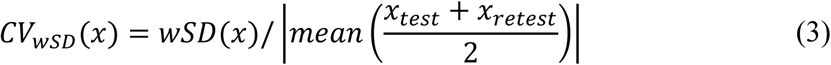

Similarly, CVBias was calculated as:

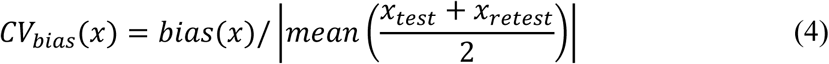

CVwSD and CVBias were expressed as percentages.

Statistical comparisons were performed using paired non-parametric Wilcoxon signed-rank tests implemented with the MATLAB built-in function, ‘signrank’. Pairwise comparisons were performed across acquisition types and, where applicable, across amplitude-based measures. For each paired comparison, the effect size was calculated from the normal approximation statistic (Z) returned by ‘signrank’:

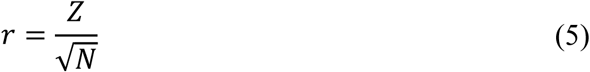

where N is the number of paired observations included in the comparison. Bonferroni correction was applied within each family of pairwise comparisons and corrected (p < 0.05) was considered statistically significant.

### 3.5. Echo combination effect on cortical coverage and reliability

The effect of T_2_*-weighted echo combination and reliability on cortical coverage was examined by comparing the multi-echo combined and the middle-echo datasets. For each participant, a cortical coverage mask was generated by binarizing the base EPI image using ‘3dAutomask’ in AFNI and projecting the resulting mask onto the ‘suma_TT_N27′ surface model in SUMA (Saad, Reynolds et al. 2004). After projection, a vertex-wise cortical coverage map was generated by calculating, at each surface vertex, the percentage of participants whose EPI mask covered that vertex.

The effect of echo combination on ICC was examined on all 400 cortical parcels. For each ROI, FC reliability was summarized as the median ICC across all 399 edges connected to that ROI, and fALFF reliability was assessed using the ICC of the ROI-wise fALFF value. To quantify the relative difference in reliability between the two data sets, the absolute percent difference (APD) in ICC between MEPI and MMID was then calculated for each ROI as:

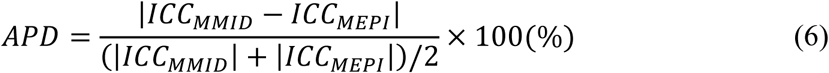

ROI-wise APD values were summarized by measure and network.

## 4. Results

### 4.1. fMRI dataset description

The dataset included 22 test–retest pairs of multi-echo multiband fMRI and 21 test–retest pairs of single-echo multiband fMRI, with a mean test–retest interval of 15.7 ± 4.0 days. Because dataset pairs from two volunteers were excluded due to excessive motion, the final analysis included 20 pairs for MEPI and MMID and 19 pairs for SEPI. Figure 1 summarizes temporal SNR and motion across sessions for each acquisition type. No significant session differences were observed in temporal SNR or motion derivative, indicating that the included datasets had comparable session-level data quality. Paired questionnaire data were available for 19 participants. The self-reported measures were generally comparable between sessions, whereas acquisition time was modestly earlier during Session 2, with a median within-participant difference of 0.35 hours (Supplementary Figure 1).

**Figure 1.**
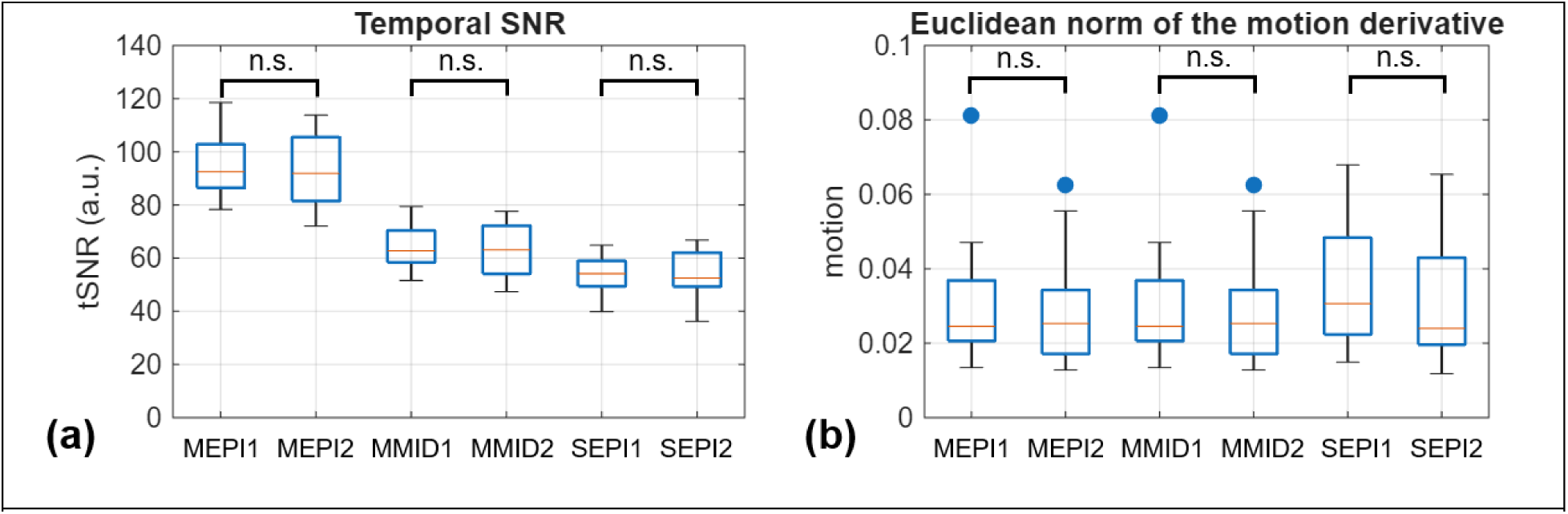
Temporal SNR and head motion summary of the resting-state fMRI datasets used for test–retest analysis. Boxplots show (a) temporal SNR and (b) Euclidean norm of the motion derivative for each acquisition and session. For each acquisition type, the suffixes 1 and 2 indicate the first (test) and second (retest) sessions, respectively (MEPI1/MEPI2, MMID1/MMID2, and SEPI1/SEPI2). Brackets indicate paired session comparisons within each acquisition type using the Wilcoxon signed-rank test. No significant session differences were observed for either temporal SNR or motion.

### 4.2. Reliability of Functional Connectivity

Functional connectivity was calculated from the Schaefer 400 cortical parcels, resulting in 79,800 unique edges. For each edge, FC strength and test–retest reliability metrics were evaluated. Parcels belonging to the Limbic A and Limbic B networks, which contain 13 and 11 parcels, respectively, were excluded from comparative analysis because many of these parcels exhibited substantial susceptibility-related signal dropout. After excluding edges involving these networks, 4,980 within-network edges and 65,520 between-network edges were included in the analysis.

Table 2 summarizes within-network FC strength and reliability metrics across acquisition types. Median within-network FC strength was highest for MEPI, followed by SEPI and MMID. In contrast, median ICC values were highly comparable across acquisitions, indicating similar reliability. Absolute within-subject variability, measured by wSD, were low for all acquisitions, with SEPI showing slightly lower wSD values than MEPI and MMID. After normalization by FC strength, CVwSD was similar across acquisitions and corresponded to approximately 28.9–31.4% of FC strength. Systematic test–retest bias was also small across all acquisition types CVBias corresponded to approximately 5.7–6.7% of FC strength.

**Table 2.**
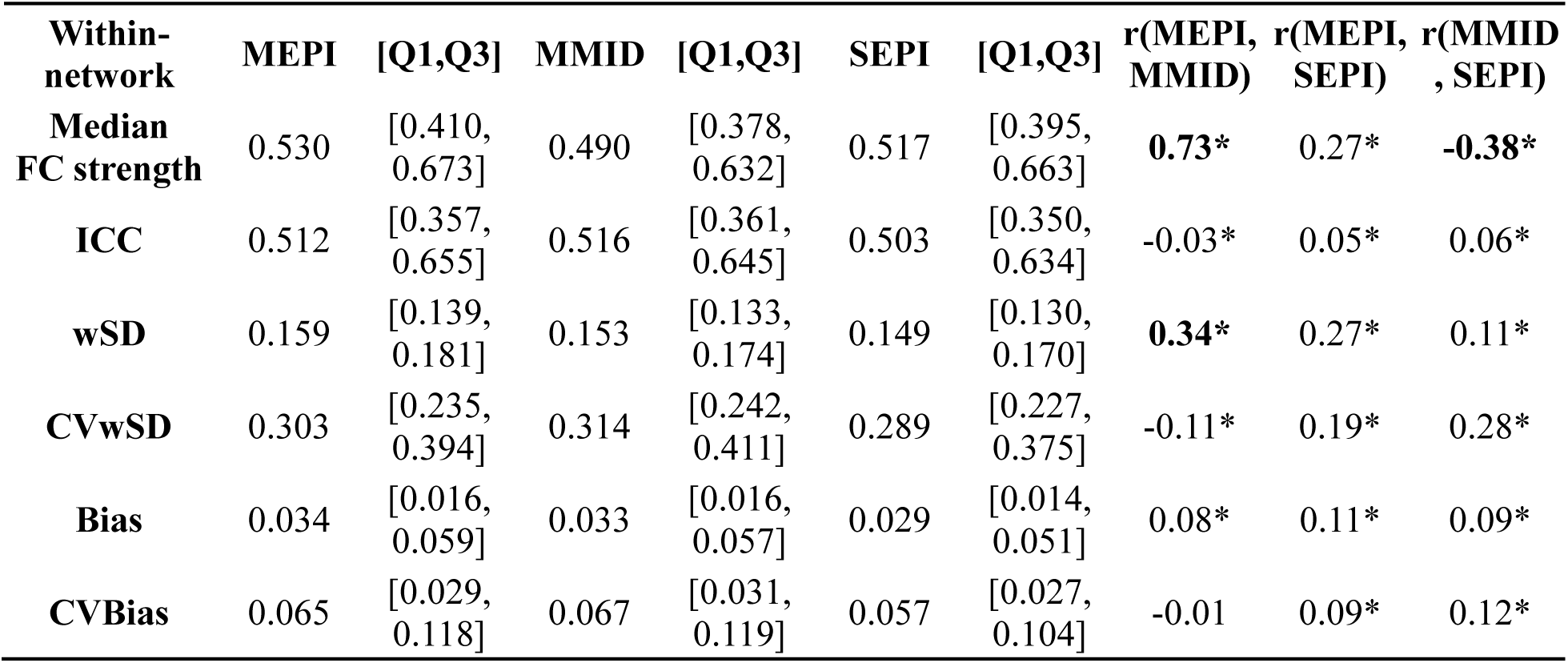
Within-network Edge-wise FC strength and test–retest reliability across acquisitions.

| Within-network | MEPI | [Q1,Q3] | MMID | [Q1,Q3] | SEPI | [Q1,Q3] | r(MEPI, MMID) | r(MEPI, SEPI) | r(MMID, SEPI) |
| --- | --- | --- | --- | --- | --- | --- | --- | --- | --- |
| **Median FC strength** | 0.530 | [0.410, 0.673] | 0.490 | [0.378, 0.632] | 0.517 | [0.395, 0.663] | **0.73\*** | 0.27* | **-0.38\*** |
| **ICC** | 0.512 | [0.357, 0.655] | 0.516 | [0.361, 0.645] | 0.503 | [0.350, 0.634] | -0.03* | 0.05* | 0.06* |
| **wSD** | 0.159 | [0.139, 0.181] | 0.153 | [0.133, 0.174] | 0.149 | [0.130, 0.170] | **0.34\*** | 0.27* | 0.11* |
| **CVwSD** | 0.303 | [0.235, 0.394] | 0.314 | [0.242, 0.411] | 0.289 | [0.227, 0.375] | -0.11* | 0.19* | 0.28* |
| **Bias** | 0.034 | [0.016, 0.059] | 0.033 | [0.016, 0.057] | 0.029 | [0.014, 0.051] | 0.08* | 0.11* | 0.09* |
| **CVBias** | 0.065 | [0.029, 0.118] | 0.067 | [0.031, 0.119] | 0.057 | [0.027, 0.104] | -0.01 | 0.09* | 0.12* |

Table 3 summarizes the corresponding results for between-network FC edges. Similar to the within-network findings, FC strength, ICC, and bias were comparable across acquisition types. The wSD and CVwSD values tended to be slightly higher for MEPI than for MMID and SEPI, although the associated effect sizes were generally small. Compared with the within-network FC, between-network edges showed lower median FC strength and modestly lower ICC values. Because wSD values were of similar magnitude to those observed for within-network edges, the lower FC strength resulted in substantially higher CVwSD and CVBias values for between-network edges.

**Table 3.** Between-network Edge-wise FC strength and test–retest reliability across acquisitions.

| Between-network | MEPI | [Q1,Q3] | MMID | [Q1,Q3] | SEPI | [Q1,Q3] | r(MEPI, MMID) | r(MEPI, SEPI) | r(MMID, SEPI) |
| --- | --- | --- | --- | --- | --- | --- | --- | --- | --- |
| **Median FC strength** | 0.211 | [0.171, 0.289] | 0.211 | [0.170, 0.287] | 0.209 | [0.164, 0.291] | 0.09* | 0.09* | 0.02* |
| **ICC** | 0.443 | [0.289, 0.578] | 0.437 | [0.289, 0.572] | 0.442 | [0.292, 0.576] | 0.03* | 0.00 | -0.01* |
| **wSD** | 0.152 | [0.134, 0.171] | 0.146 | [0.130, 0.164] | 0.138 | [0.122, 0.156] | **0.33\*** | **0.36\*** | 0.22* |
| **CVwSD** | 0.985 | [0.580, 2.011] | 0.891 | [0.550, 1.671] | 0.816 | [0.507, 1.501] | **0.34\*** | **0.35\*** | 0.20* |
| **Bias** | 0.032 | [0.015, 0.055] | 0.032 | [0.015, 0.055] | 0.029 | [0.013, 0.049] | 0.00 | 0.09* | 0.10* |
| **CVBias** | 0.206 | [0.085, 0.493] | 0.196 | [0.083, 0.435] | 0.167 | [0.070, 0.381] | 0.11* | 0.16* | 0.11* |

Overall, acquisition-dependent difference in FC reliability was present but limited. Although several pairwise signed-rank comparisons showed statistically significant acquisition effects for several metrics, the corresponding effect sizes for reliability metrics were generally small. In contrast, the distinction between within-and between-network FC exerted a larger influence on reliability characteristics, with between-network connections exhibiting lower FC strength and ICC.

#### 4.2.1. Network-wise reliability of functional connectivity

We next examined whether reliability varied according to large-scale functional network organization. Figure 2 summarizes network-wise FC strength and reliability metrics by grouping edges according to the 15-network Schaefer classification after exclusion of the Limbic A and B networks. Each matrix cell represents the median value across edges belonging to the corresponding network pair. Diagonal elements represent within-network edges, whereas off-diagonal elements represent between-network edges. In Figure 2a, median FC strength showed a pronounced network structure, with substantial higher values for within-network than between-network edges. The strongest within-network FC was observed in the Somatomotor A, central and peripheral Visual, and Dorsal Attention A networks. However, the corresponding ICC matrix (Figure 2b) showed a different spatial pattern. Higher ICC values were concentrated in network pairs involving the Control and Default Mode networks, whereas somatomotor and visual network-related pairs exhibited only moderate reliability despite relatively strong FC strength.

**Figure 2.**
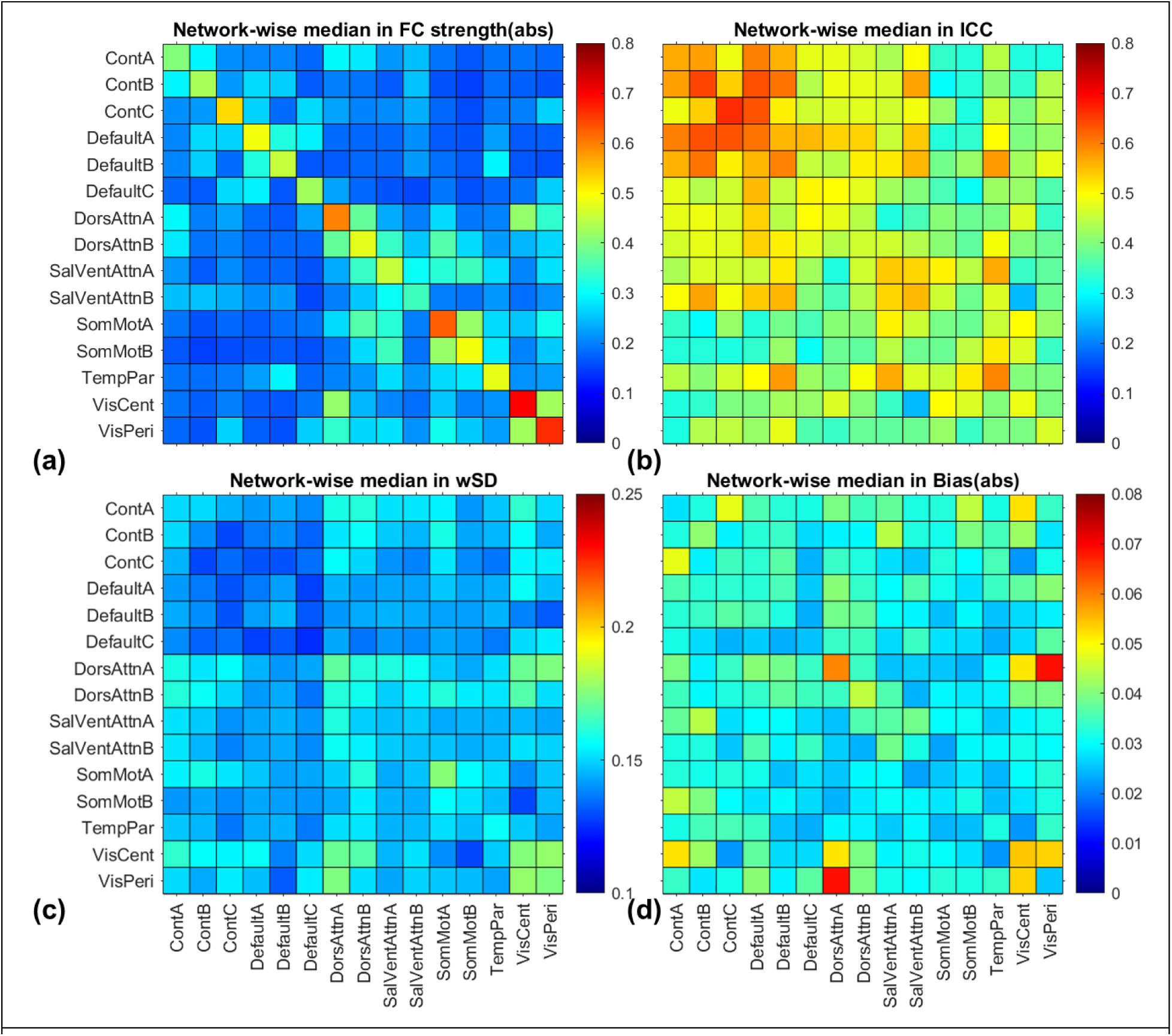
Network-wise matrices for (a) functional connectivity strength and test-retest reliability metrics of (b) ICC, (c) wSD, and (d) systematic bias. Each cell represents median values of edgewise metrics. These results were obtained with median values across MEPI, MMID, and SEPI datasets.

A similar pattern was observed in the wSD matrix in Figure 2c, where network pairs involving the default mode and control networks tended to show lower wSD values. In contrast, network pairs with lower ICC generally showed higher wSD values. The bias matrix (Figure 2d) demonstrated uniformly low values across most network pairs, with only a few localized regions exhibiting slightly elevated bias. Overall, these findings indicate that FC reliability is strongly influenced by large-scale functional network organization and is not solely determined by the magnitude of functional connectivity.

Figure 3 further integrates the complementary reliability information provided by ICC and wSD. Figure 3a shows a scatter plot of network-wise median FC values in the ICC–wSD space. Percentile thresholds for ICC and wSD were used to classify network pairs into four reliability categories, with higher ICC and lower wSD indicating more favorable reliability. Among 120 network-wise FC cells, 10 were classified into the most reliable category, followed by 26, 36, and 48 cells in the remaining categories. To visualize the spatial distribution of these reliability categories, the classification results were mapped onto the network-wise FC matrix in Figure 3b. The most reliable network pairs, shown in red, were mainly observed in network pairs involving Default A, with additional reliable pairs involving Control B and C. In contrast, networks with generally high FC strength, such as Somatomotor A and Visual Central networks, were not classified among the most reliable network-wise pairs. These results indicate that FC magnitude alone did not determine test–retest reliability.

**Figure 3.**
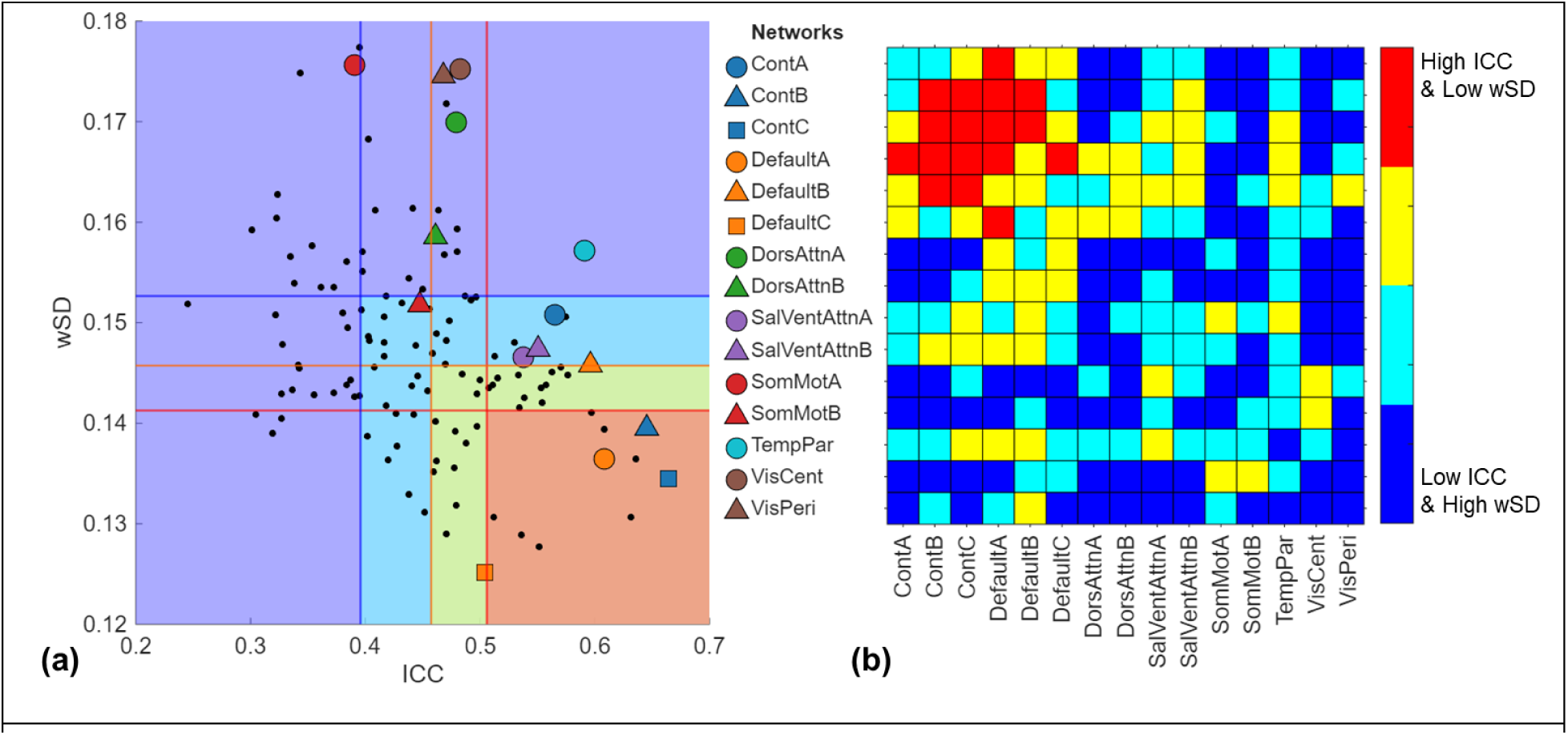
Network-wise classification of FC reliability based on ICC and wSD. (a) Scatter plot of network-wise median ICC and wSD values. Within-network FC values are indicated by network-specific markers, and between-network FC values are shown as black dots. Vertical and horizontal lines indicate the 25th, 50th, and 75th percentile thresholds for ICC and wSD, respectively. Reliability categories were defined based on higher ICC and lower wSD, with red indicating the most reliable category, followed by yellow, light blue, and dark blue. (b) The reliability categories were mapped onto the network-wise FC matrix to visualize their spatial distribution.

### 4.3. Reliability of amplitude-based resting-state fMRI measures

Among amplitude-based resting-state fMRI measures, ALFF is commonly used to quantify the magnitude of spontaneous low-frequency BOLD signal fluctuations, whereas fALFF and mALFF represent normalized variants of this measure. To determine which amplitude-based measure showed the most favorable mid-term test–retest reliability, ICC, CVwSD, and CVBias were evaluated for ALFF, fALFF, and mALFF. Because these measures have different numerical scales, scale-normalized variability and bias metrics were used for comparisons across measures. Consistent with the FC analysis, parcels belonging to the Limbic A and Limbic B networks were excluded, resulting in 376 ROIs for this analysis.

Figure 4 summarizes the reliability profiles of ALFF, fALFF, and mALFF across MEPI, MMID, and SEPI acquisitions. Figure 4a shows ICC values for each measure and acquisition type. Pairwise comparisons across measures were performed using ROI-level median values across the three acquisitions. ICC values were broadly comparable across ALFF, fALFF, and mALFF, with no clear measure-dependent difference. In contrast, clearer differences emerged for the scale-normalized measures of variability and bias. fALFF consistently exhibited lower CVwSD than both ALFF and mALFF across all acquisitions (Figure 4b), indicating reduced within-subject variability relative to signal magnitude. Similarly, fALFF demonstrated lower CVBias values than ALFF and mALFF (Figure 4c), reflecting reduced systematic test–retest shifts.

**Figure 4.**
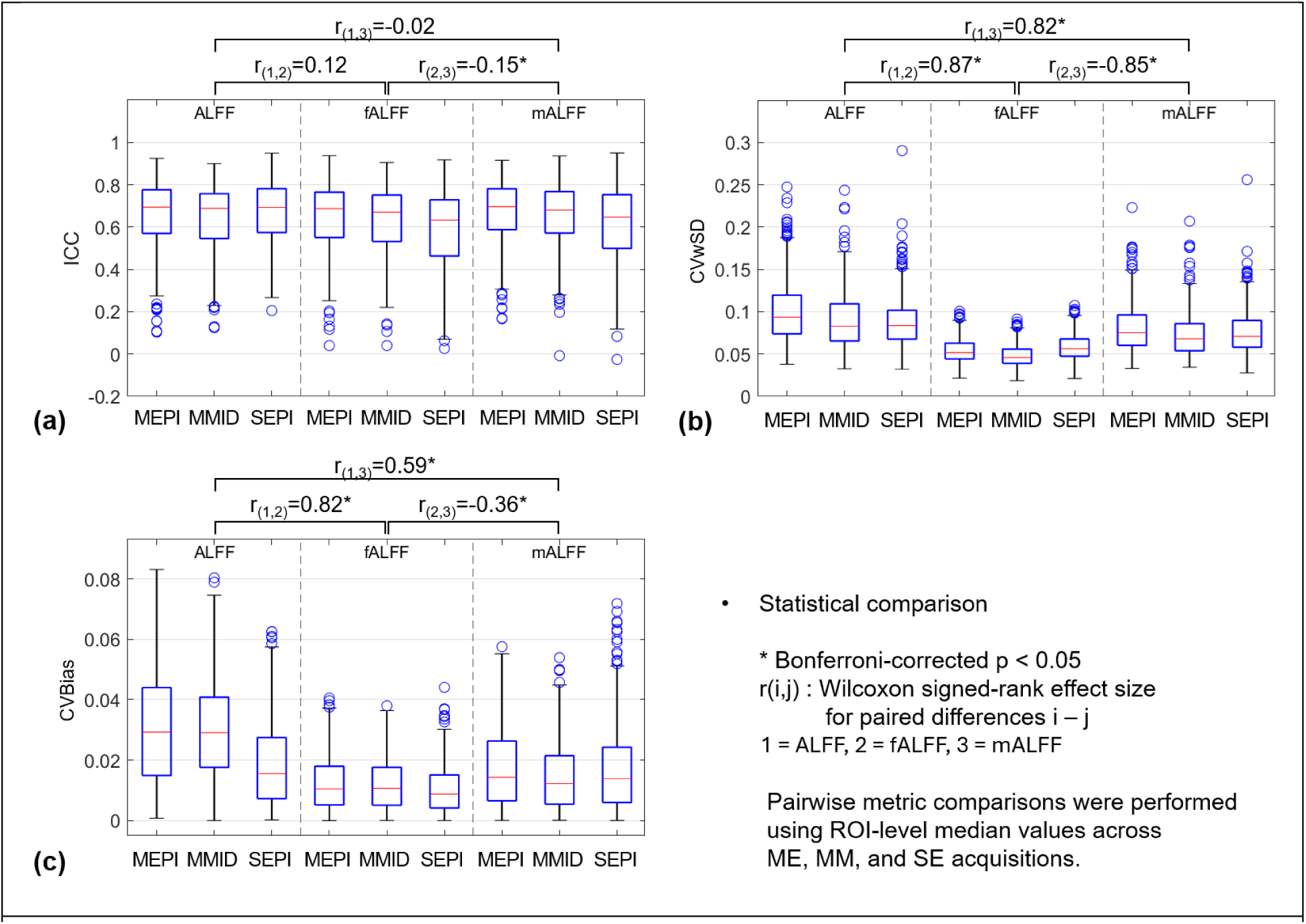
Reliability of amplitude-based resting-state fMRI measures across acquisitions. Boxplots show ROI-level distributions of (a) ICC, (b) CVwSD, and (c) CVBias for ALFF, fALFF, and mALFF measured from ME, MM, and SE acquisitions. Pairwise comparisons among ALFF, fALFF, and mALFF were performed using ROI-level median values across the three acquisitions. Brackets indicate Wilcoxon signed-rank effect sizes for paired metric differences, where 1 = ALFF, 2 = fALFF, and 3 = mALFF. Asterisks indicate Bonferroni-corrected significance at p < 0.05.

Notably, the differences among ALFF, fALFF, and mALFF were larger and more consistent than the corresponding differences among MEPI, MMID, and SEPI. Within each amplitude measures, the three acquisition strategies showed highly similar ICC, CVwSD, and CVBias distributions. In contrast, fALFF provided the most favorable reliability profile among the three amplitude-based resting-state fMRI measures, particularly in terms of relative within-subject variability and systematic bias. These findings suggest that a selection of amplitude-based resting-state fMRI measures has a greater influence on test–retest stability than acquisition strategy on the Compact 3T platform. Because fALFF consistently combined high ICC with low relative variability and bias, subsequent amplitude-based analyses focused on fALFF.

Table 4 summarizes ROI-wise fALFF magnitude and test-retest reliability metrics across acquisition types. Median fALFF magnitude was highest for MEPI, followed by MMID and SEPI, indicating that acquisition strategy influenced the measured signal amplitude. ICC values followed a similar order, with MEPI showing slightly higher reliability than MMID and SEPI, although the corresponding pairwise effect sizes were small. All three acquisitions exhibited comparable reliability profiles. Although MEPI tended to have slightly higher wSD values, CVwSD remained below 6% for all acquisition types. Systematic test–retest bias was minimal, with CVBias values of approximately 1% across acquisitions. Overall, these findings indicate that acquisition strategy had a measurable effect on fALFF magnitude but only a limited influence on its test–retest reliability.

**Table 4.** ROI-wise fALFF magnitude and test–retest reliability across acquisitions.

|  | MEPI | [Q1,Q3] | MMID | [Q1,Q3] | SEPI | [Q1,Q3] | r(MEPI, MMID) | r(MEPI, SEPI) | r(MMID, SEPI) |
| --- | --- | --- | --- | --- | --- | --- | --- | --- | --- |
| **Median fALFF** | 0.273 | [0.247, 0.299] | 0.246 | [0.227, 0.267] | 0.193 | [0.175, 0.210] | **0.87\*** | **0.87\*** | **0.87\*** |
| **ICC** | 0.687 | [0.551, 0.765] | 0.670 | [0.532, 0.751] | 0.633 | [0.463, 0.729] | 0.24* | 0.34* | 0.23* |
| **wSD** | 0.014 | [0.012, 0.018] | 0.012 | [0.009, 0.015] | 0.011 | [0.009, 0.014] | **0.86\*** | **0.79\*** | 0.22* |
| **CVwSD** | 0.052 | [0.044, 0.063] | 0.046 | [0.039, 0.056] | 0.056 | [0.047, 0.068] | **0.76\*** | -0.34* | **-0.74\*** |
| **Bias** | 0.003 | [0.001, 0.005] | 0.003 | [0.001, 0.004] | 0.002 | [0.001, 0.003] | 0.21* | 0.43* | 0.36* |
| **CVBias** | 0.010 | [0.005, 0.018] | 0.011 | [0.005, 0.018] | 0.009 | [0.004, 0.015] | 0.01 | 0.17* | 0.17* |

Compared with the FC reliability results in Tables 2 and 3, fALFF showed substantially greater measurement stability. Across acquisitions, median fALFF ICC values ranged from 0.633 to 0.687, exceeding those observed for within-network FC. Likewise, median fALFF CVwSD values ranged from 4.6% to 5.6%, markedly lower than the corresponding FC values. Although regional amplitude measures and edge-wise FC characterize different aspects of resting-state brain function, these results suggest that spontaneous fluctuation amplitude is more reproducible than functional connectivity over a 2–3 week interval. Taken together with the results in Figure 4, the findings indicate that the choice of resting-state measure exerts a larger influence on reliability than the choice of EPI acquisition on the Compact 3T platform.

#### 4.3.1. Network-wise reliability of fALFF measure

To determine whether the network-dependent reliability patterns observed for FC were also present in amplitude-based measures, fALFF reliability was evaluated across 15 networks after excluding the Limbic A and Limbic B networks. Figure 5 summarizes the distributions of fALFF magnitude and reliability metrics across ROIs within each network. Figure 5a shows the distribution of fALFF magnitude, while Figures 5b–d show the corresponding distributions of ICC, wSD, and bias, respectively. The dotted horizontal line indicates the global median across all included ROIs for each metric.

**Figure 5.**
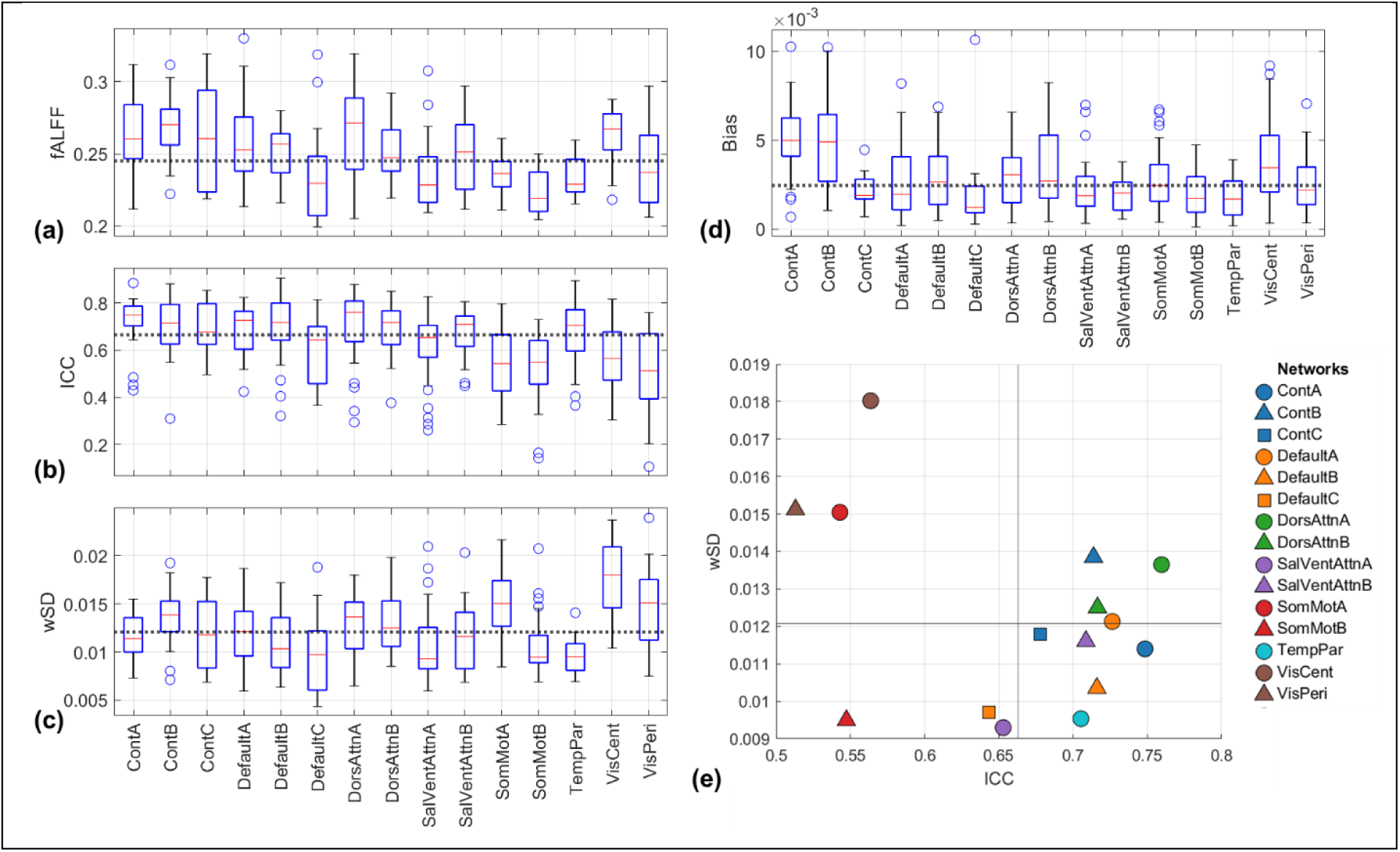
Network-wise distribution of fALFF magnitude and test–retest reliability. Boxplots show ROI-level distributions of (a) fALFF magnitude, (b) ICC, (c) wSD, and (d) bias across 15 networks after excluding the Limbic A and Limbic B networks. The dotted horizontal line indicates the global median across all included ROIs for each metric. (e) Scatter plot of network-wise median ICC and wSD values. Each point represents one network, with marker color and shape indicating network identity. Vertical and horizontal reference lines indicate the global median ICC and wSD, respectively.

Network-dependent differences in fALFF reliability were evident, although the range of variation was smaller than that observed for FC. Several association networks, including the default mode, control, attention, and temporoparietal networks, showed favorable reliability profiles characterized with relatively high ICC and low wSD values. In contrast, somatomotor and visual networks showed less favorable reliability profiles, driven by lower ICC, higher wSD, or both. Bias values were low across all networks, without a clear network-specific pattern.

Figure 5e summarizes the joint relationship between network-wise ICC and wSD using network median values. Most networks were clustered within a relatively narrow range, indicating broadly comparable fALFF reliability across networks. Networks located in the lower-right quadrant combined high ICC with low within-subject variability and therefore exhibited the most favorable reliability profiles. These networks were predominantly control, dorsal attention, salience/ventral attention, and temporoparietal systems. Conversely, somatomotor and visual networks tended to occupy regions with lower ICC or higher wSD values. Among association networks, some networks with relatively high ICC also exhibited moderately elevated wSD, whereas others combined high ICC with low wSD. Thus, relative and absolute reliability metrics did not consistently identify the same networks as most reliable. These results suggest that fALFF reliability was generally consistent across networks, with particularly favorable reliability observed in temporoparietal and selected association networks, and modestly reduced reliability in several sensory and motor networks.

### 4.4. Effect of susceptibility-related geometric distortion on reliability

The spatial relationship between susceptibility-related geometric distortion and EPI signal dropout versus test–retest reliability was examined with a particular focus on inferior frontal and temporal regions that are prone to susceptibility-related signal loss. Figure 6a shows the cortical distribution of Schaefer 400 parcels grouped into Yeo’s 17 networks on lateral and medial views of the SUMA surface template. Because EPI signal dropout is commonly observed in inferior frontal and temporal regions, Figure 6b shows inferior surface views of the network distribution together with EPI coverage maps for MEPI and MMID. EPI coverage was higher for MEPI than for MMID in several inferior cortical regions, including regions partially overlapping with Limbic A and B.

**Figure 6.**
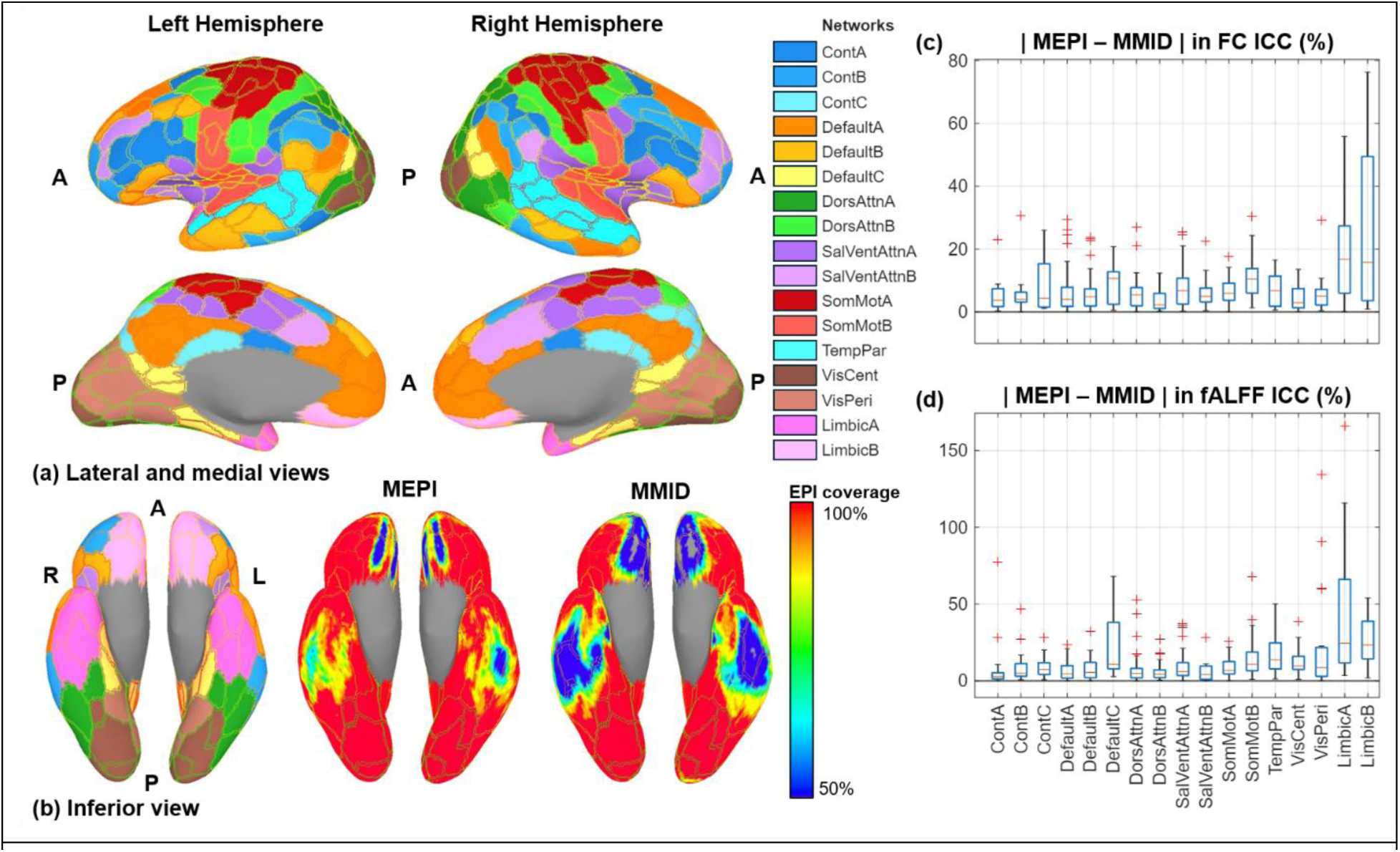
Geometric distortion-related spatial heterogeneity of reliability in regions prone to susceptibility-related EPI signal dropout. (a) Lateral and medial surface views of the SUMA surface template with cortical network labels. (b) Inferior surface views showing network labels and EPI coverage for MEPI and MMID. EPI coverage was defined as the percentage of participants with valid EPI signal coverage at each surface location after preprocessing. Inferior frontal and temporal regions are prone to susceptibility-related signal dropout in EPI and partially overlap with parcels in the Limbic A and B networks. (c, d) ROI-wise absolute percentage differences in ICC between MEPI and MMID for within-network FC and fALFF, summarized by network. Limbic A and B showed larger and more variable MEPI– MMID ICC differences than most other networks, indicating greater spatial heterogeneity in reliability differences in susceptibility-prone inferior cortical regions.

To assess whether these spatial differences were reflected in reliability, we calculated ROI-wise absolute percentage differences in ICC between MEPI and MMID and summarized them by network. Figure 6c shows the MEPI–MMID ICC differences for FC reliability, and Figure 6d shows the corresponding differences for fALFF reliability. Limbic A and B showed larger and more variable MEPI–MMID ICC differences than most other networks. These findings indicate that MEPI–MMID reliability differences in susceptibility-prone inferior frontal and temporal regions were spatially heterogeneous rather than uniform. The spatial pattern suggests that acquisition-dependent differences in EPI coverage may contribute to reliability differences in these regions.

### 4.5. Effect of Multi-echo ICA-based denoising

Multi-echo ICA-based denoising produced distinct rsfMRI measures and reliability profiles compared with the conventional preprocessing pipelines, including MEPI, MMID, and SEPI. As shown in Table 5, MICA yielded median absolute FC strengths of 0.997 for within-network FC and 0.655 for between-network FC, both substantially higher than those obtained with MEPI. MICA yielded a higher median ICC than MEPI for between-network FC but a slightly lower median ICC for within-network FC. In contrast, median wSD was higher for MICA than for MEPI for both within-network FC and between-network FC indicating increased absolute scan-to-scan variability. However, because MICA also increased FC strength, the corresponding median CVwSD values were lower than those of MEPI. Absolute test–retest bias was also larger for MICA than for MEPI. However, CVBias was modest for within-network FC and was lower for between-network FC because of the increased FC magnitude.

**Table 5.** ME-ICA-based values and test–retest reliability across fMRI measures.

| **MICA** | **Within-**<br>**network**<br>**FC** | **[Q1,Q3]** | **r(MEPI,**<br>**MICA)** | **Between-**<br>**network**<br>**FC** | **[Q1,Q3]** | **r(MEPI, MICA)** | **ROI-wise**<br>**fALFF** | **[Q1,Q3]** | **r(MEPI,**<br>**MICA)** |
| --- | --- | --- | --- | --- | --- | --- | --- | --- | --- |
| **Value** | 0.997 | [0.845,<br>1.147] | -0.87* | 0.655 | [0.544,<br>0.777] | -0.87* | 0.310 | [0.285,<br>0.337] | -0.87* |
| **ICC** | 0.465 | [0.336,<br>0.582] | 0.23* | 0.515 | [0.419,<br>0.598] | -0.35* | 0.491 | [0.345,<br>0.609] | 0.82* |
| **wSD** | 0.222 | [0.195,<br>0.249] | -0.85* | 0.215 | [0.194,<br>0.237] | -0.86* | 0.024 | [0.021,<br>0.028] | -0.87* |
| **CVwSD** | 0.223 | [0.189,<br>0.263] | 0.82* | 0.326 | [0.269,<br>0.405] | 0.85* | 0.078 | [0.065,<br>0.093] | -0.86* |
| **Bias** | 0.068 | [0.036,<br>0.102] | -0.49* | 0.064 | [0.034,<br>0.096] | -0.52* | 0.010 | [0.007,<br>0.012] | -0.84* |
| **CVBias** | 0.069 | [0.036,<br>0.103] | 0.06* | 0.097 | [0.051,<br>0.152] | 0.60* | 0.032 | [0.024,<br>0.040] | -0.83* |

r(MEPI, MICA): effect size for MEPI vs. MICA. Positive values indicate MEPI > MICA; negative values indicate MICA > MEPI. MEPI values for within-network FC, between-network FC, and ROI-wise fALFF are shown in Tables 2, 3, and 4, respectively.

For fALFF, MICA showed a different pattern from FC. Compared with MEPI, MICA yielded higher absolute fALFF amplitude, wSD, CVwSD, bias, and CVBias, but lower ICC. Thus, fALFF derived from MICA appeared less reliable than fALFF derived from MEPI across the evaluated reliability metrics.

Figure 7 shows the network-pair FC strength and reliability profiles for MICA. Absolute FC strength was generally high across network pairs, with the largest values observed for within-network connections, particularly within the SomMotA, SomMotB, VisCent, and VisPeri networks, as well as for several connections among the somatomotor, salience/ventral-attention, and visual networks.. Network-pair ICC values in MICA were relatively high for many connections involving the Control and Default Mode subnetworks, whereas lower ICC values were observed for several connections involving SomMotB and the dorsal-attention and salience/ventral-attention networks..

**Figure 7.**
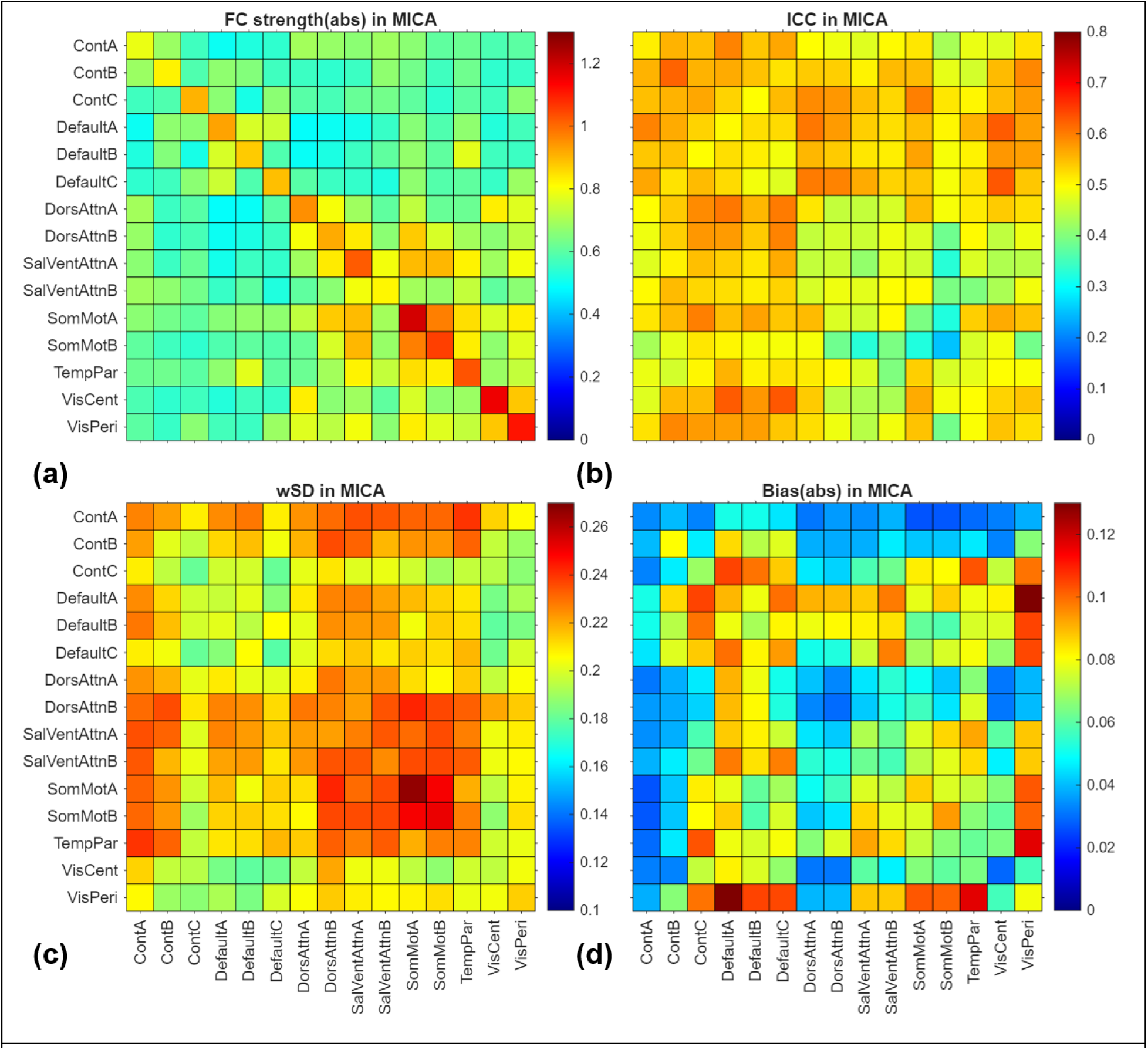
Network-wise matrices for (a) functional connectivity strength and test-retest reliability metrics of (b) ICC, (c) wSD, and (d) systematic bias. These results were obtained with MICA datasets.

Network-pair wSD and bias in MICA were generally higher than those observed in MEPI (Figure 2), but also showed distinct network-dependent patterns. For wSD, lower values were observed in several network pairs involving Default Mode subnetworks and Visual networks. For bias, lower values were observed in several network pairs involving ContA, ContB, DorsAttnA, and DorsAttnB. Overall, MICA produced a reliability pattern that differed from that of MEPI, indicating that multi-echo ICA-based denoising substantially reshaped downstream FC estimates and their reliability characteristics rather than uniformly improving or worsening test–retest reliability.

## 5. Discussion

This study characterized the mid-term test–retest reliability of resting-state fMRI measures across multi-echo, a middle-echo dataset derived from the multi-echo acquisition and independently acquired single-echo multiband EPI acquisitions on a compact 3T MRI system with high-performance gradients. The main findings were as follows. First, acquisition-dependent differences in reliability were present but generally modest. MEPI produced higher FC and fALFF magnitudes in several comparisons, but these signal-magnitude differences did not consistently translate into lower within-subject variability (wSD) or substantially higher ICC. Second, FC reliability showed network-pair-specific patterns and was not determined solely by FC strength (Wisner, Atluri et al. 2013). Third, among amplitude-based resting-state fMRI measures, fALFF showed the most favorable overall reliability profile, with ICC comparable to those of ALFF and mALFF but substantially lower wSD and bias. Fourth, ROI-wise fALFF showed higher ICC and markedly lower wSD than edge-wise FC in this dataset, although this comparison should be interpreted cautiously because the two measures have different data structures, biological interpretations, and noise characteristics.

A central question in this study was whether multi-echo acquisition provides a test–retest reliability advantage over middle-echo–derived or conventional single-echo imaging on the compact 3T platform. The results regarding FC measures and reliability demonstrate an important distinction between signal magnitude and measurement reliability: an acquisition may increase the measured FC value without necessarily improving its stability across sessions. Since FC is sensitive to head motion and physiological or brain-state fluctuations (Power, Barnes et al. 2012), the comparison between MEPI and MMID is particularly informative. The similar reliability of MEPI and MMID suggests that optimal echo combination altered FC magnitude more than it altered mid-term reproducibility under the processing strategy used in this study. The comparable performance of SEPI further indicates that the additional acquisition and processing complexity of MEPI did not yield a broad reliability advantage for FC on this platform.

For fALFF, the acquisition effect was somewhat more favorable to MEPI. The principal effect of MEPI on fALFF was an increase in measured fluctuation amplitude, accompanied by only a modest improvement in relative reliability (i.e. ICC). Higher fALFF magnitude may reflect increased sensitivity to low-frequency BOLD fluctuations, but magnitude alone cannot establish improved biological validity or sensitivity to disease-or intervention-related effects. Taken together, the FC and fALFF results indicate that the benefits of multi-echo acquisition were metric-dependent and more apparent for amplitude magnitude than for test–retest reliability.

The distinction between within-network and between-network FC had a larger influence on reliability characteristics than acquisition type. Within-network edges showed substantially higher FC strength and modestly higher ICC than between-network edges. In contrast, absolute within-subject variability was broadly comparable between the two edge classes. The higher ICC of within-network FC may therefore partly reflect greater between-subject variance and stronger signal magnitude rather than substantially lower scan-to-scan variability. Note that scale-normalized metrics such as CVwSD and CVBias should be interpreted cautiously for FC edges or network pairs with low or near-zero FC magnitude in between-network FC. Nevertheless, the results demonstrate that longitudinal or intervention-related FC changes should be evaluated relative to the expected variability of the specific network pair rather than against a single whole-brain reliability threshold.

The use of ICC, wSD, and bias also revealed information that would have been missed by ICC alone. Some network pairs showed relatively high ICC despite moderate wSD, whereas others had low wSD but only moderate ICC. Such differences are expected because ICC reflects the ratio of between-subject variance to total variance and may therefore be relatively high despite substantial scan-to-scan variability when between-subject differences are large. Accordingly, ICC and wSD should not be expected to rank acquisition strategies, functional networks, or preprocessing pipelines identically. Scale-normalized measures such as CVwSD and CVBias also require careful interpretation because they are influenced by the magnitude of the underlying rsfMRI measure, particularly when FC values are low or near zero. These findings emphasize the importance of jointly considering relative reliability, absolute variability, systematic bias, and signal magnitude rather than relying on a single reliability metric. Systematic bias was generally low and showed little network-specific organization, suggesting that differences in reliability were driven primarily by random within-subject variability and preservation of inter-individual differences rather than by a consistent shift between test and retest sessions.

Among ALFF, fALFF, and mALFF, fALFF showed the most favorable reliability profile. ICC values were broadly comparable across the three measures, with mALFF showing slightly higher ICC (Jia, Sun et al. 2020). In contrast, fALFF exhibited substantially lower CVwSD and CVBias than ALFF and mALFF across all acquisition types. The normalization of low-frequency fluctuation amplitude by the broader frequency spectrum may reduce the influence of global signal scaling, scanner drift, and nonspecific fluctuations, thereby improving relative within-subject stability. However, because this normalization may also alter between-subject variance and physiological sensitivity, the lower CVwSD of fALFF should not be interpreted as evidence that fALFF is universally more sensitive to neural, disease-related, or intervention-related effects. Rather, ALFF and fALFF may reflect related but distinct physiological aspects of spontaneous BOLD fluctuations (Aiello, Salvatore et al. 2015, Deng, Franklin et al. 2022), and their reliability profiles should be interpreted accordingly.

The complementary reliability metrics used in this study also allowed comparison between amplitude-based measures and FC. Compared with edge-wise FC, ROI-wise fALFF showed higher ICC and substantially lower relative within-subject variability, suggesting that regional amplitude-based measures may be more stable than edge-wise connectivity measures. However, this comparison should be interpreted cautiously because FC and fALFF have different data structures and noise characteristics. Moreover, recent studies suggest that FC and amplitude-based measures may capture different aspects of spontaneous brain activity and provide complementary information. Therefore, it may be important to understand their distinct characteristics and limitations and to interpret them together in longitudinal or intervention studies, rather than selecting one measure over the other. Also, within the acquisition conditions evaluated here, differences among resting-state fMRI measures were larger and more consistent than differences among acquisition strategies. These variability estimates provide an empirical reference for evaluating whether longitudinal or intervention-related changes exceed expected scan-to-scan variability. They may also inform power calculations and sample size planning for future studies using specific resting-state fMRI metrics or network pairs.

Network-wise analyses showed reliability was spatially heterogeneous and metric-specific. For FC, favorable combinations of high ICC and low wSD were frequently observed in network pairs involving Default A and selected control networks. By contrast, several Somatomotor and Visual network pairs exhibited strong FC magnitude but were not among the most reliable. This dissociation reinforces the conclusion that stronger connectivity does not necessarily imply greater temporal stability. Intersession variability in these networks may also partly reflect differences in behavioral state. Because participants were scanned with their eyes open, subtle changes in visual attention or fixation across sessions could have affected Visual network measures, whereas small differences in posture or body movement not fully captured by head-motion estimates could have influenced Somatomotor network measures. For fALFF, reliability was broadly consistent across networks, but selected association networks, including the default mode, control, attention, and temporoparietal networks, showed favorable reliability profiles (Blautzik, Keeser et al. 2013, Wisner, Atluri et al. 2013, Zuo and Xing 2014). These network-wise patterns were broadly consistent with prior observations that higher reliability can be found in default mode and frontoparietal/control networks.

In the present study, the visual and somatomotor networks showed less favorable reliability for both FC and fALFF consistent with some previous reports (Chen, Xu et al. 2015, Somandepalli, Kelly et al. 2015). However, this pattern differs from other studies reporting relatively high reliability in these networks (Blautzik, Keeser et al. 2013, Wisner, Atluri et al. 2013). Such discrepancies may arise from differences in nuisance-regression strategies, including physiological-noise correction, motion regression, and global signal regression, as well as differences in parcellation scheme, sample characteristics, and reliability metric (Shirer, Jiang et al. 2015, Noble, Scheinost et al. 2019). The method used to estimate FC may be particularly important (Shirer, Jiang et al. 2015). Some prior studies evaluated ICA-or dual-regression-based intrinsic connectivity networks (Blautzik, Keeser et al. 2013, Wisner, Atluri et al. 2013), whereas the present study used atlas-defined ROI-to-ROI pairwise FC edges. In addition, studies relying exclusively on ICC may rank networks differently from studies that jointly consider ICC and absolute variability. The network categories used here were defined relative to the distributions observed in this dataset and should therefore be viewed as descriptive rather than universal reliability thresholds.

These results provide several practical considerations for studies performed on compact high-performance MRI systems. First, resting-state measurements obtained with MEPI, MMID, and SEPI showed broadly comparable mid-term reliability, indicating that robust longitudinal measurements can be obtained with each acquisition strategy. Second, increased signal or metric magnitude should not be assumed to represent increased reproducibility. Third, expected variability depends on the metric and network under study. Consequently, an observed longitudinal change should ideally be evaluated against reliability estimates matched to the acquisition, metric, and functional system being examined.

The approximately 2-3 week interval used here captures both measurement variability and natural changes in participant state across independent sessions. The reported reliability values should therefore not be interpreted as pure measures of technical repeatability or immutable trait-like organization. Rather, they represent the combined stability of the acquisition, processing pipeline, physiological state, and spontaneous brain activity over a clinically relevant mid-term interval. This makes them particularly relevant as non-intervention benchmarks for determining whether changes following stimulation, treatment, disease progression, or recovery exceed expected scan-to-scan and day-to-day variation.

Multi-echo combination may be particularly relevant in cortical regions affected by susceptibility-related EPI signal loss. In this study, MEPI showed greater cortical coverage than MMID in portions of the inferior frontal and temporal cortex. Networks overlapping these susceptibility-prone regions, particularly Limbic A and B networks (Somandepalli, Kelly et al. 2015), showed relatively large and variable MEPI–MMID differences in ICC. This pattern suggests that the reliability effect of multi-echo combination may be most apparent in regions where signal loss and signal recovery are spatially prominent.

Although this study primarily characterized the reliability of T2*-weighted combined multi-echo data, we also included a supplementary benchmark using multi-echo independent component analysis with the default tedana workflow (Kundu, Inati et al. 2012, DuPre, Salo et al. 2021). Previous work has suggested that ME-ICA-based processing may improve reliability when assessed by the similarity of spatial FC patterns (Lynch, Power et al. 2020). Using complementary test-retest metrics, including ICC, wSD, and test-retest bias, we found that MICA substantially altered downstream FC estimates rather than uniformly improving or worsening reliability. For FC, MICA substantially increased absolute FC magnitude and improved between-network ICC, while slightly reducing within-network ICC and increasing wSD and absolute bias. However, the corresponding normalized variability and bias measures were lower or remained modest, partly because of the larger FC magnitude. In contrast, MICA-derived fALFF showed lower ICC and higher absolute and normalized variability and bias than MEPI-derived fALFF, indicating less favorable reliability across the evaluated metrics. These results suggest that MICA produces a distinct reliability profile and highlight that preprocessing choices may meaningfully alter the magnitude and test–retest characteristics of multi-echo fMRI.

Several limitations should be noted. First, the sample size was modest, which may reduce the precision of ICC estimates and the stability of network-level rankings. Only healthy volunteers were included, and reliability may differ in older adults, patients, or populations with greater motion and physiological variability. Second, this was a single-site study performed on a compact 3T MRI system with high-performance gradient coils, which may limit generalizability across sites, scanner platforms, coil configurations, or image-reconstruction pipelines. No direct comparison with a conventional whole-body 3T scanner was performed, so the findings should not be interpreted as demonstrating superior reliability of the compact platform relative to conventional systems. Third, although the present study did not evaluate all possible preprocessing strategies, the differences observed between conventional MEPI processing and ME-ICA denoising suggest that reliability estimates can be meaningfully influenced by processing choices. This underscores the need to benchmark preprocessing pipelines when test– retest reliability is a primary consideration. Fourth, MEPI and SEPI differed in TR and the number of acquired volumes. These differences were inherent to the sequence designs and may have influenced temporal sampling and measurement precision as well as subject state and motion. MMID was derived from the MEPI acquisition and therefore provided a controlled comparison of echo combination. Nevertheless, by characterizing mid-term test–retest reliability in the absence of intervention, this study provides an estimate of trait-like stability under healthy conditions. These reliability estimates may serve as a non-intervention benchmark for determining whether disease-related, stimulation-induced, or treatment-related changes exceed expected measurement and day-to-day variability.

In conclusion, this study provides test–retest reliability benchmarks for resting-state fMRI measures acquired with MEPI, MMID, and SEPI on a compact 3T MRI system. Acquisition-dependent reliability was generally modest. MEPI increased FC strength and fALFF magnitude in several comparisons, but these increases did not consistently translate into substantially improved reliability. FC reliability showed network-pair-specific patterns that were not simply determined by FC strength. Among amplitude-based measures, fALFF showed the most favorable combination of comparable ICC, low relative within-subject variability, and low bias. Collectively, these findings indicate that metric and network selection may influence measurement stability as much as, or more than, echo-sampling strategy and provide practical reference values for interpreting longitudinal and intervention-related resting-state fMRI changes on the compact 3T.

## 7. Ethics approval statement

The study was conducted in accordance with the Declaration of Helsinki, and approved by the Institutional Review Board of Mayo Clinic (IRB 14-001036). And written informed consent was obtained from all participants.

## 8. Acknowledgements

The authors thank Tim L. Waters and Erin M. Gray for their assistance with study coordination, and Maria A. Halverson, Chloe A. Pallex, and Derek G. Bischof for their technical assistance with magnetic resonance imaging data acquisition. This work was supported by National Institute of Biomedical Imaging and Bioengineering (NIBIB) at the National Institutes of Health (NIH).

## 9. Author contributions

**Daehun Kang**: Conceptualization, Methodology, Formal analysis, Writing - Original Draft, Visualization, **Kirk M. Welker:** Methodology, **Dora Hermes**: Methodology, **Matt A. Bernstein:** Methodology, **John Huston III:** Methodology**, Yunhong Shu:** Conceptualization, Methodology, Writing - Original Draft, **All**: Writing - Review & Editing

## 12. Data availability statement

The datasets generated and analyzed in this study are not publicly available due to institutional regulations and data protection policies governing protected health information.

## 13. Declaration of generative AI and AI-assisted technologies in the writing process

During the preparation of this work the author used ChatGPT powered by GPT-5.6 Sol (OpenAI, accessed July 21, 2026) in order to improve language and readability. After using the ChatGPT powered by GPT-5.6 Sol, the authors reviewed and edited the content as needed and take full responsibility for the content of the publication.

## References

1. Aiello, M., E. Salvatore, A. Cachia, S. Pappatà, C. Cavaliere, A. Prinster, E. Nicolai, M. Salvatore, J. C. Baron and M. Quarantelli (2015). “Relationship between simultaneously acquired resting-state regional cerebral glucose metabolism and functional MRI: A PET/MR hybrid scanner study.” Neuroimage 113: 111–121.

2. Birn, R. M., M. A. Smith, T. B. Jones and P. A. Bandettini (2008). “The respiration response function: the temporal dynamics of fMRI signal fluctuations related to changes in respiration.” Neuroimage 40(2): 644–654.

3. Biswal, B., F. Z. Yetkin, V. M. Haughton and J. S. Hyde (1995). “Functional connectivity in the motor cortex of resting human brain using echo-planar MRI.” Magn Reson Med 34(4): 537–541.

4. Bland, J. M. and D. G. Altman (1986). “Statistical methods for assessing agreement between two methods of clinical measurement.” Lancet 1(8476): 307–310.

5. Blautzik, J., D. Keeser, A. Berman, M. Paolini, V. Kirsch, S. Mueller, U. Coates, M. Reiser, S. J. Teipel and T. Meindl (2013). “Long-term test-retest reliability of resting-state networks in healthy elderly subjects and with amnestic mild cognitive impairment patients.” J Alzheimers Dis 34(3): 741–754.

6. Cahart, M. S., O. O’Daly, V. Giampietro, M. Timmers, J. Streffer, S. Einstein, F. Zelaya, F. Dell’Acqua and S. C. R. Williams (2023). “Comparing the test-retest reliability of resting-state functional magnetic resonance imaging metrics across single band and multiband acquisitions in the context of healthy aging.” Hum Brain Mapp 44(5): 1901–1912.

7. Chen, B., T. Xu, C. Zhou, L. Wang, N. Yang, Z. Wang, H. M. Dong, Z. Yang, Y. F. Zang, X. N. Zuo and X. C. Weng (2015). “Individual Variability and Test-Retest Reliability Revealed by Ten Repeated Resting-State Brain Scans over One Month.” PLoS One 10(12): e0144963.

8. Cohen, A. D., B. Yang, B. Fernandez, S. Banerjee and Y. Wang (2021). “Improved resting state functional connectivity sensitivity and reproducibility using a multiband multi-echo acquisition.” Neuroimage 225: 117461.

9. Conwell, K., B. von Reutern, N. Richter, J. Kukolja, G. R. Fink and O. A. Onur (2018). “Test-retest variability of resting-state networks in healthy aging and prodromal Alzheimer’s disease.” Neuroimage Clin 19: 948–962.

10. Cox, R. W. (1996). “AFNI: software for analysis and visualization of functional magnetic resonance neuroimages.” Comput Biomed Res 29(3): 162–173.

11. Deng, S., C. G. Franklin, M. O’Boyle, W. Zhang, B. L. Heyl, P. A. Jerabek, H. Lu and P. T. Fox (2022). “Hemodynamic and metabolic correspondence of resting-state voxel-based physiological metrics in healthy adults.” Neuroimage 250: 118923.

12. DuPre, E., T. Salo, Z. Ahmed, P. Bandettini, K. Bottenhorn, C. Caballero-Gaudes, L. Dowdle, J. Gonzalez-Castillo, S. Heunis, P. Kundu, A. Laird, R. Markello, C. Markiewicz, S. Moia, I. Staden, J. Teves, E. Uruñuela, M. Vaziri-Pashkam, K. Whitaker and D. Handwerker (2021). “TE-dependent analysis of multi-echo fMRI with tedana.” Journal of Open Source Software 6(66).

13. Feinberg, D. A., A. J. S. Beckett, A. T. Vu, J. Stockmann, L. Huber, S. Ma, S. Ahn, K. Setsompop, X. Cao, S. Park, C. Liu, L. L. Wald, J. R. Polimeni, A. Mareyam, B. Gruber, R. Stirnberg, C. Liao, E. Yacoub, M. Davids, P. Bell, E. Rummert, M. Koehler, A. Potthast, I. Gonzalez-Insua, S. Stocker, S. Gunamony and P. Dietz (2023). “Next-generation MRI scanner designed for ultra-high-resolution human brain imaging at 7 Tesla.” Nat Methods 20(12): 2048– 2057.

14. Finn, E. S., X. Shen, D. Scheinost, M. D. Rosenberg, J. Huang, M. M. Chun, X. Papademetris and R. Todd Constable (2015). “Functional connectome fingerprinting: identifying individuals using patterns of brain connectivity.” Nature Neuroscience 18(October): 1–11.

15. Foo, T. K. F., E. Laskaris, M. Vermilyea, M. Xu, P. Thompson, G. Conte, C. Van Epps, C. Immer, S. K. Lee, E. T. Tan, D. Graziani, J. B. Mathieu, C. J. Hardy, J. F. Schenck, E. Fiveland, W. Stautner, J. Ricci, J. Piel, K. Park, Y. Hua, Y. Bai, A. Kagan, D. Stanley, P. T. Weavers, E. Gray, Y. Shu, M. A. Frick, N. G. Campeau, J. Trzasko, J. Huston, 3rd and M. A. Bernstein (2018). “Lightweight, compact, and high-performance 3T MR system for imaging the brain and extremities.” Magn Reson Med 80(5): 2232–2245.

16. Foo, T. K. F., E. T. Tan, M. E. Vermilyea, Y. H. Hua, E. W. Fiveland, J. E. Piel, K. Park, J. Ricci, P. S. Thompson, D. Graziani, G. Conte, A. Kagan, Y. Bai, C. Vasil, M. Tarasek, D. T. B. Yeo, F. Snell, D. Lee, A. Dean, J. K. DeMarco, R. Y. Shih, M. N. Hood, H. Chae and V. B. Ho (2020). “Highly efficient head-only magnetic field insert gradient coil for achieving simultaneous high gradient amplitude and slew rate at 3.0T (MAGNUS) for brain microstructure imaging.” Magnetic Resonance in Medicine 83(6): 2356–2369.

17. Geerligs, L., M. Rubinov, R. N. Henson and Cam-CAN (2015). “State and Trait Components of Functional Connectivity: Individual Differences Vary with Mental State.” Journal of Neuroscience 35(41): 13949–13961.

18. Glover, G. H., T. Q. Li and D. Ress (2000). “Image-based method for retrospective correction of physiological motion effects in fMRI: RETROICOR.” Magnetic Resonance in Medicine 44(1): 162–167.

19. Gorgolewski, K. J., N. Mendes, D. Wilfling, E. Wladimirow, C. J. Gauthier, T. Bonnen, F. J. Ruby, R. Trampel, P. L. Bazin, R. Cozatl, J. Smallwood and D. S. Margulies (2015). “A high resolution 7-Tesla resting-state fMRI test-retest dataset with cognitive and physiological measures.” Sci Data 2: 140054.

20. Heunis, S., M. Breeuwer, C. Caballero-Gaudes, L. Hellrung, W. Huijbers, J. F. Jansen, R. Lamerichs, S. Zinger and A. P. Aldenkamp (2021). “The effects of multi-echo fMRI combination and rapid T2*-mapping on offline and real-time BOLD sensitivity.” Neuroimage 238: 118244.

21. Jia, X. Z., J. W. Sun, G. J. Ji, W. Liao, Y. T. Lv, J. Wang, Z. Wang, H. Zhang, D. Q. Liu and Y. F. Zang (2020). “Percent amplitude of fluctuation: A simple measure for resting-state fMRI signal at single voxel level.” PLoS One 15(1): e0227021.

22. Jo, H. J., S. J. Gotts, R. C. Reynolds, P. A. Bandettini, A. Martin, R. W. Cox and Z. S. Saad (2013). “Effective Preprocessing Procedures Virtually Eliminate Distance-Dependent Motion Artifacts in Resting State FMRI.” J Appl Math 2013.

23. Jo, H. J., Z. S. Saad, W. K. Simmons, L. A. Milbury and R. W. Cox (2010). “Mapping sources of correlation in resting state FMRI, with artifact detection and removal.” Neuroimage 52(2): 571– 582.

24. Kang, D., M. H. In, H. J. Jo, M. A. Halverson, N. K. Meyer, Z. Ahmed, E. M. Gray, R. Madhavan, T. K. Foo, B. Fernandez, D. F. Black, K. M. Welker, J. D. Trzasko, J. Huston, 3rd, M. A. Bernstein and Y. Shu (2023). “Improved Resting-State Functional MRI Using Multi-Echo Echo-Planar Imaging on a Compact 3T MRI Scanner with High-Performance Gradients.” Sensors (Basel) 23(9).

25. Kang, D., H. J. Jo, M. H. In, U. Yarach, N. K. Meyer, L. J. Bardwell Speltz, E. M. Gray, J. D. Trzasko, J. Huston Iii, M. A. Bernstein and Y. Shu (2020). “The benefit of high-performance gradients on echo planar imaging for BOLD-based resting-state functional MRI.” Phys Med Biol 65(23): 235024.

26. Kang, D., K. Uchida, C. R. Haider, N. G. Campeau, M. H. In, E. M. Gray, J. D. Trzasko, K. M. Welker, M. A. Bernstein, M. R. Trenerry, D. R. Holmes Iii, M. J. Joyner, T. B. Curry, J. Huston Iii and Y. Shu (2026). “Brain functional connectivity initiates structured reorganization at a critical oxygen threshold during hypoxia.” Brain Res Bull 235: 111748.

27. Kundu, P., S. J. Inati, J. W. Evans, W.-M. M. Luh and P. A. Bandettini (2012). “Differentiating BOLD and non-BOLD signals in fMRI time series using multi-echo EPI.” NeuroImage 60(3): 1759–1770.

28. Lynch, C. J., J. D. Power, M. A. Scult, M. Dubin, F. M. Gunning and C. Liston (2020). “Rapid Precision Functional Mapping of Individuals Using Multi-Echo fMRI.” Cell Reports 33(12): 108540.

29. McGraw, K. O. and S. P. Wong (1996). “Forming inferences about some intraclass correlation coefficients.” Psychological Methods 1(1): 30–46.

30. Mejia, A. F., M. B. Nebel, H. Shou, C. M. Crainiceanu, J. J. Pekar, S. Mostofsky, B. Caffo and M. A. Lindquist (2015). “Improving reliability of subject-level resting-state fMRI parcellation with shrinkage estimators.” Neuroimage 112: 14–29.

31. Ning, L., N. Makris, J. A. Camprodon and Y. Rathi (2019). “Limits and reproducibility of resting-state functional MRI definition of DLPFC targets for neuromodulation.” Brain Stimul 12(1): 129–138.

32. Noble, S., D. Scheinost and R. T. Constable (2019). “A decade of test-retest reliability of functional connectivity: A systematic review and meta-analysis.” Neuroimage 203: 116157.

33. Noble, S., M. N. Spann, F. Tokoglu, X. Shen, R. T. Constable and D. Scheinost (2017). “Influences on the Test-Retest Reliability of Functional Connectivity MRI and its Relationship with Behavioral Utility.” Cereb Cortex 27(11): 5415–5429.

34. O’Connor, D., N. V. Potler, M. Kovacs, T. Xu, L. Ai, J. Pellman, T. Vanderwal, L. C. Parra, S. Cohen, S. Ghosh, J. Escalera, N. Grant-Villegas, Y. Osman, A. Bui, R. C. Craddock and M. P. Milham (2017). “The Healthy Brain Network Serial Scanning Initiative: a resource for evaluating inter-individual differences and their reliabilities across scan conditions and sessions.” Gigascience 6(2): 1–14.

35. Pannunzi, M., R. Hindriks, R. G. Bettinardi, E. Wenger, N. Lisofsky, J. Martensson, O. Butler, E. Filevich, M. Becker, M. Lochstet, S. Kuhn and G. Deco (2017). “Resting-state fMRI correlations: From link-wise unreliability to whole brain stability.” Neuroimage 157: 250–262.

36. Poser, B. a., M. J. Versluis, J. M. Hoogduin and D. G. Norris (2006). “BOLD contrast sensitivity enhancement and artifact reduction with multiecho EPI: Parallel-acquired inhomogeneity-desensitized fMRI.” Magnetic Resonance in Medicine 55(6): 1227–1235.

37. Posse, S., S. Wiese, D. Gembris, K. Mathiak, C. Kessler, M. L. Grosse-Ruyken, B. Elghahwagi, T. Richards, S. R. Dager and V. G. Kiselev (1999). “Enhancement of BOLD-contrast sensitivity by single-shot multi-echo functional MR imaging.” Magn. Reson. Med. 42(1): 87–97.

38. Power, J. D., K. A. Barnes, A. Z. Snyder, B. L. Schlaggar and S. E. Petersen (2012). “Spurious but systematic correlations in functional connectivity MRI networks arise from subject motion.” Neuroimage 59(3): 2142–2154.

39. Ramos-Llorden, G., H. H. Lee, M. Davids, P. Dietz, A. Krug, J. E. Kirsch, M. Mahmutovic, A. Muller, Y. Ma, H. Lee, C. Maffei, A. Yendiki, B. Bilgic, D. J. Park, Q. Tian, B. Clifford, W. C. Lo, S. Stocker, J. Fischer, G. Ruyters, M. Roesler, A. Potthast, T. Benner, E. Rummert, R. Schuster, P. J. Basser, T. Witzel, L. L. Wald, B. R. Rosen, B. Keil and S. Y. Huang (2026). “Ultra-high gradient connectomics and microstructure MRI scanner for imaging of human brain circuits across scales.” Nat Biomed Eng 10(2): 309–324.

40. Reuter, M., N. J. Schmansky, H. D. Rosas and B. Fischl (2012). “Within-subject template estimation for unbiased longitudinal image analysis.” Neuroimage 61(4): 1402–1418.

41. Saad, Z. S., R. C. Reynolds, B. Argall, S. Japee and R. W. Cox (2004). SUMA: An interface for surface-based intra-and inter-subject analysis with AFNI. 2004 2nd IEEE International Symposium on Biomedical Imaging: Macro to Nano. 2: 1510–1513.

42. Salarian, A. (2016). Intraclass Correlation Coefficient (ICC), MATLAB Central File Exchange.

43. Schaefer, A., R. Kong, E. M. Gordon, T. O. Laumann, X. N. Zuo, A. J. Holmes, S. B. Eickhoff and B. T. T. Yeo (2018). “Local-Global Parcellation of the Human Cerebral Cortex from Intrinsic Functional Connectivity MRI.” Cereb Cortex 28(9): 3095–3114.

44. Setsompop, K., B. A. Gagoski, J. R. Polimeni, T. Witzel, V. J. Wedeen and L. L. Wald (2012). “Blipped-controlled aliasing in parallel imaging for simultaneous multislice echo planar imaging with reduced g-factor penalty.” Magn Reson Med 67(5): 1210–1224.

45. Shirer, W. R., H. Jiang, C. M. Price, B. Ng and M. D. Greicius (2015). “Optimization of rs-fMRI Pre-processing for Enhanced Signal-Noise Separation, Test-Retest Reliability, and Group Discrimination.” Neuroimage 117: 67–79.

46. Somandepalli, K., C. Kelly, P. T. Reiss, X. N. Zuo, R. C. Craddock, C. G. Yan, E. Petkova, F. X. Castellanos, M. P. Milham and A. Di Martino (2015). “Short-term test-retest reliability of resting state fMRI metrics in children with and without attention-deficit/hyperactivity disorder.” Dev Cogn Neurosci 15: 83–93.

47. Wisner, K. M., G. Atluri, K. O. Lim and A. W. Macdonald, 3rd (2013). “Neurometrics of intrinsic connectivity networks at rest using fMRI: retest reliability and cross-validation using a meta-level method.” Neuroimage 76: 236–251.

48. Wu, A., J. Ricci, M. Xu, V. Soni, G. Conte, C. Van Epps, M. Parizh, W. Stautner, Y. Hua, S. K. Lee, M. Vermilyea, D. T. Yeo and T. K. Foo (2026). “Cooldown and Ramp Test of a Low-Cryogen, Lightweight, Head-Only 7T MRI Magnet.” IEEE Trans Appl Supercond 36(3).

49. Yang, H., X. Y. Long, Y. H. Yang, H. Yan, C. Z. Zhu, X. P. Zhou, Y. F. Zang and Q. Y. Gong (2007). “Amplitude of low frequency fluctuation within visual areas revealed by resting-state functional MRI.” Neuroimage 36(1): 144–152.

50. Zang, Y. F., Y. He, C. Z. Zhu, Q. J. Cao, M. Q. Sui, M. Liang, L. X. Tian, T. Z. Jiang and Y. F. Wang (2007). “Altered baseline brain activity in children with ADHD revealed by resting-state functional MRI.” Brain Dev 29(2): 83–91.

51. Zou, Q. H., C. Z. Zhu, Y. Yang, X. N. Zuo, X. Y. Long, Q. J. Cao, Y. F. Wang and Y. F. Zang (2008). “An improved approach to detection of amplitude of low-frequency fluctuation (ALFF) for resting-state fMRI: fractional ALFF.” J Neurosci Methods 172(1): 137–141.

52. Zuo, X. N., A. Di Martino, C. Kelly, Z. E. Shehzad, D. G. Gee, D. F. Klein, F. X. Castellanos, B. B. Biswal and M. P. Milham (2010). “The oscillating brain: complex and reliable.” Neuroimage 49(2): 1432–1445.

53. Zuo, X. N., C. Kelly, J. S. Adelstein, D. F. Klein, F. X. Castellanos and M. P. Milham (2010). “Reliable intrinsic connectivity networks: test-retest evaluation using ICA and dual regression approach.” Neuroimage 49(3): 2163–2177.

54. Zuo, X. N. and X. X. Xing (2014). “Test-retest reliabilities of resting-state FMRI measurements in human brain functional connectomics: a systems neuroscience perspective.” Neurosci Biobehav Rev 45: 100–118.

