## Supplemental Figure 1 for "Network- and Measure-Specific Mid-Term Reliability of Multi-Echo Resting-State Functional Magnetic Resonance Imaging on a Compact 3 Tesla Scanner"

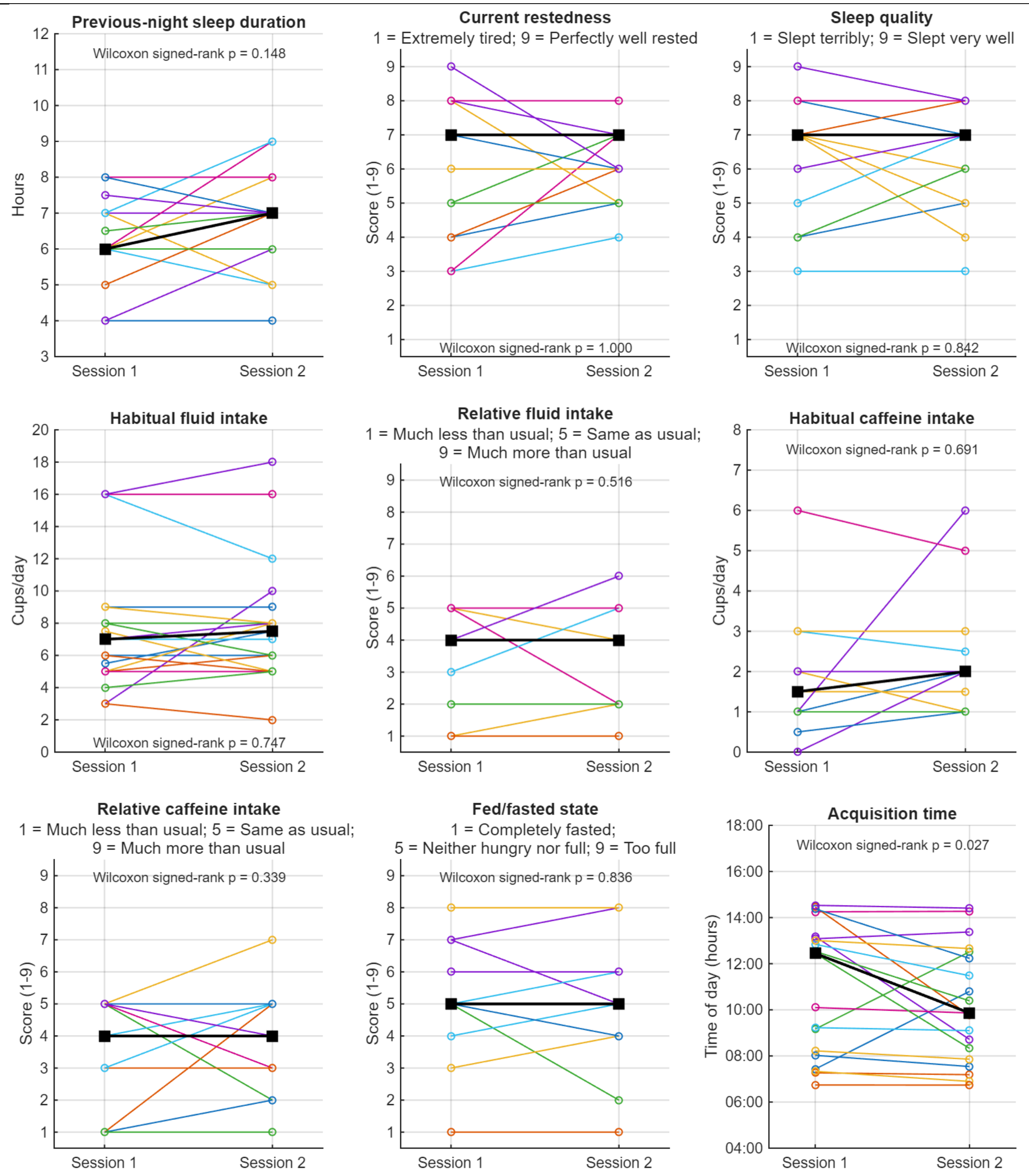


Supplementary Figure 1. **Session-to-session comparison of behavioral, physiological, and acquisition-related measures.** Colored lines and open circles represent individual participants, whereas black squares connected by black lines indicate the median for each session. Previous-night sleep duration, current restedness, sleep quality, habitual and relative fluid intake, habitual and relative caffeine intake, fed/fasted state, and scan acquisition time were compared between Sessions 1 and 2 using Wilcoxon signed-rank tests. The full questionnaire and response scales are provided in Supplementary Material 1. No significant between-session differences were observed in the self-reported measures, whereas acquisition time was modestly earlier during Session 2. Because several questionnaire variables were measured using discrete ordinal scales, some individual observations and trajectories overlap.
