## Supplemental Material 1 for "Network- and Measure-Specific Mid-Term Reliability of Multi-Echo Resting-State Functional Magnetic Resonance Imaging on a Compact 3 Tesla Scanner"

### Behavioral and Physiological Questionnaire

1. Scan date: ______________________
2. Which hand is dominant? Circle one: Left or Right
3. How many times have you been scanned on the Compact 3T? ________
4. Do you have any physical discomfort today? No _____ Yes _____ (If yes, please explain below.)

   _________________________________________________________________________
5. On average, how many hours do you sleep every night? ___________________ hours
6. How many hours did you sleep last night? ___________________ hours
7. How well rested do you feel right now?

| Extremely tired | | | | | | | | Perfectly well rested |
|---|---|---|---|---|---|---|---|---|
| 1 | 2 | 3 | 4 | 5 | 6 | 7 | 8 | 9 |

8. How well did you sleep last night?

| I slept terribly | | | | | | | | I slept very well |
|---|---|---|---|---|---|---|---|---|
| 1 | 2 | 3 | 4 | 5 | 6 | 7 | 8 | 9 |

9. On average, how much water (and other liquids) do you drink every day (in cups[1])? __________
10. Comparing to other days did you drink more or less water today?

| much less than usual | | | | the same amount as usual | | | | much more than usual |
|---|---|---|---|---|---|---|---|---|
| 1 | 2 | 3 | 4 | 5 | 6 | 7 | 8 | 9 |

11. On average, how much caffeinated drinks (coffee, caffeinated soda[2], non-herbal tea[3], etc.) do you drink every day (in cups[1])? __________________
12. Comparing to other days did you drink more or less coffee and other caffeinated drinks (coffee, caffeinated soda[2], non-herbal tea[3], etc.) today?

| much less than usual | | | | the same amount as usual | | | | much more than usual |
|---|---|---|---|---|---|---|---|---|
| 1 | 2 | 3 | 4 | 5 | 6 | 7 | 8 | 9 |

13. How was your fed or fasted state just before participating in this scan?

| Completely fasted | | | | Neither hungry nor full | | | | Too full |
|---|---|---|---|---|---|---|---|---|
| 1 | 2 | 3 | 4 | 5 | 6 | 7 | 8 | 9 |

**Thank you for taking the time to complete this survey.**

**[1] 8 ounces = 1 cup, 5 cups ~ 1 liter**
**[2] e.g. Cola, Mountain Dew, Sunkist, etc.**
**[3] e.g. Black Tea, Green Tea, Oolong Tea, Pu-erh Tea, White Tea, etc.**
